# Symbiotic State Affects Microbiome Recovery in a Facultatively Symbiotic Cnidarian

**DOI:** 10.1101/2025.11.04.686571

**Authors:** Maria Valadez-Ingersoll, Caoimhe A. Bodnar, Emily X. Feng, Audrey Wong, Thomas D. Gilmore, Sarah W. Davies

## Abstract

The cnidarian holobiont consists of host cells, algal symbionts, and a complex microbiome that resides in and on the host tissue and the algal symbionts. To investigate the interactions between a host cnidarian, its algal symbionts, and its bacterial microbiome, we used antibiotics to deplete the microbiome of the facultatively symbiotic sea anemone *Exaiptasia pallida* (Aiptasia) in both symbiotic and aposymbiotic states. To understand the dynamics of microbiome reestablishment following disruption, we used 16S gene sequencing of the bacterial microbiome to characterize communities during recovery following antibiotic treatment. We assessed the host molecular response to microbiome depletion and recovery using RNA-seq and Western blotting of the immune transcription factor NF-κB. 16S results demonstrate that, following depletion, symbiotic Aiptasia can reestablish bacterial communities (based on diversity metrics and community composition) alike to those of controls before microbiome depletion. However, the microbiomes of aposymbiotic Aiptasia fail to reestablish to control conditions even after seven days of recovery. Bacteria in the Family Endozoicomonadaceae reestablish to control levels in symbiotic, but not aposymbiotic, Aiptasia, suggesting an association between Endozoicomonadaceae and algal symbionts. Our molecular analyses show that host immune system gene expression is downregulated during recovery from antibiotic treatment, but that NF-κB protein levels increase during recovery, suggesting mechanisms for microbiome reestablishment following disruption. This study shows the dynamics of microbiome recovery following depletion and the influence of microbiome community members on host gene expression in a cnidarian model, providing a foundation for future research involving pairwise interactions between microorganisms and cnidarian hosts.

## 1. Introduction

The cnidarian holobiont is composed of host cells, intracellular photosynthetic algae in the Family Symbiodiniaceae, and a microbiome comprising bacteria, archaea, viruses, fungi, and protists (Maire et al., 2022). This microbiome associates with high specificity to host microhabitats (e.g., mucus, tissue, and skeleton in calcifying hosts) (McCauley et al., 2023; Pollock et al., 2018) and algal symbionts (Ainsworth et al., 2015; Frommlet et al., 2015; Lawson et al., 2018; Maire et al., 2021). A role for the microbiome as a protective and adaptive system for cnidarian hosts in the face of environmental stress, especially pathogenic disease, has been posited by the Coral Probiotic Hypothesis (Reshef et al., 2006) and is supported by studies identifying antibiotic production by coral mucus-associated bacteria (Ritchie, 2006). The Coral Probiotic Hypothesis has been used as justification for probiotic treatment of coral with Beneficial Microorganisms for Coral (BMCs) in the laboratory and in situ (Delgadillo-Ordoñez et al., 2024; Peixoto et al., 2017). BMCs are thought to facilitate a variety of benefits to cnidarian hosts (reviewed in Peixoto et al., 2017). In particular, bacteria in the genus *Endozoicomonas* (Family Endozoicomonadaceae) have emerged as a BMC of interest, although their exact role in promoting holobiont health is still not fully understood (Peixoto et al., 2017; Pogoreutz & Ziegler, 2024). Proposed contributions of *Endozoicomonas* to holobiont health include effects on sulfur cycling and dimethylsulfoniopropionate (DMSP) transformation (Tandon et al., 2020), carbohydrate metabolism (Neave et al., 2017), and antimicrobial properties (Rua et al., 2014). Some strains of *Endozoicomonas* may participate in a tripartite interaction between the cnidarian host and their algal symbionts (Gotze, Tandon, et al., 2025).

The cnidarian microbiome can be influenced by biotic (Chan et al., 2019; Curtis et al., 2023; Röthig, Costa, et al., 2016) and abiotic (Grupstra et al., 2024; Savary et al., 2021; Ziegler et al., 2017) factors. Recent studies in several coral species have suggested a role for the bacterial microbiome in thermal resistance of the coral holobiont. In heat challenge experiments, researchers have documented a decrease in microbial taxa associated with healthy coral, including *Endozoicomonas* (Savary et al., 2021). In a meta-analysis of 45 papers documenting changes to the cnidarian microbiome in response to environmental stressors, *Endozoicomonas* abundance consistently decreased following stress events (McDevitt-Irwin et al., 2017). However, *Endozoicomonas* abundance can also decrease when corals are housed and fed in aquaria (Pogoreutz & Ziegler, 2024), highlighting the complexity of these relationships. In *Acropora hyacinthus* (Ziegler et al., 2017), *Acropora millepora*, and *Turbinaria reniformis* (Grottoli et al., 2018), corals with high tolerance to thermal bleaching harbored highly stable bacterial microbiomes. However, the protective effect of maintaining highly stable versus highly plastic microbiomes may be species-specific, as suggested by research demonstrating that slow-growing *Acropora* species altered their microbiomes in response to environmental change (termed “microbiome conformers”) whereas fast-growing *Pocillopora* species maintained more stable microbiomes under these environmental challenges (termed “microbiome regulators”) (Ziegler et al., 2019).

In addition to its potential role in protecting from thermal stress, the cnidarian microbiome has been shown to affect other factors related to holobiont health, including regeneration, disease susceptibility, and nutrient cycling. Recently, the microbiome and the host immune system have been implicated in cnidarian regeneration and wound healing (Da-Anoy et al., 2025), wherein the immune system is induced by wounding, which presumably prevents the establishment of opportunistic/pathogenic bacteria (van de Water et al., 2015). Additionally, microbiome disturbance through factors like disease and thermal stress may allow for the establishment of opportunistic or pathogenic taxa that outcompete beneficial members of the microbiome (Welsh et al., 2017; Young et al., 2023). For example, Amplicon Sequence Variants (ASVs) in the order Cytophagales increased in abundance following exposure to White Band Type I disease (Young et al., 2023) and to the coral pathogen *Vibrio coralliilyticus* (Welsh et al., 2017), which also led to an increase in the opportunistic Rhodobacterales. Opportunistic taxa often have functions associated with faster growth and higher metabolic processes, providing competitive advantages during microbiome restructuring following disturbances (Han et al., 2021; Mayali et al., 2014).

To study cnidarian host-microbial associations, efforts have been made to generate axenic (microbiome-free) cnidarians, including using antibiotics to develop gnotobiotic systems (in which the microbiome has been depleted) (Bent et al., 2021; Costa et al., 2021; MacVittie et al., 2024). The establishment of axenic cnidarians can enable investigations into the impacts of specific microbiome members on host and holobiont health, host molecular regulation of the microbiome, and the reestablishment of the microbiome following disruption (Bove et al., 2023).

To investigate the interaction between host cnidarians, their algal symbionts, and their microbiomes, we studied the dynamics of microbiome depletion and recovery in the facultatively symbiotic sea anemone *Exaiptasia pallida* (Aiptasia). We hypothesized that microbiome communities would be altered by antibiotic treatment in a symbiotic state-specific manner, but that host mechanisms for microbiome recovery would be conserved across symbiotic states. Using a transient antibiotic treatment, microbiomes were depleted in symbiotic (with algal symbionts) and aposymbiotic (without algal symbionts) Aiptasia. We then analysed the dynamics of microbiome reestablishment and the host molecular responses to microbiome depletion and recovery. While both symbiotic states downregulated gene expression pathways in the immune system during recovery, symbiotic Aiptasia had a stronger transcriptomic response than aposymbiotic Aiptasia.

Additionally, our results highlight the stability of microbiomes in symbiotic Aiptasia following disruption, which contrasts the unstable microbiomes of aposymbiotic Aiptasia following disruption. Overall, this study emphasizes the importance of an integrative approach that explores all symbiotic partners when studying aspects of the cnidarian holobiont, especially in the context of environmental changes in reef habitats.

## 2. MATERIALS AND METHODS

### 2.1 Aiptasia Maintenance and Husbandry

Aiptasia husbandry was performed as described previously (Mansfield et al., 2017; Valadez-Ingersoll et al., 2024). Briefly, adult CC7 Aiptasia in symbiosis with *Symbiodinium linucheae* were maintained in 35 ppt artificial seawater (ASW, Instant Ocean) in polycarbonate tanks at room temperature (approximately 25°C) under a 12h:12h light:dark cycle (white fluorescent light, 25 μmol photons m^-2^ sec^-1^). Water changes were performed weekly unless otherwise noted. Aiptasia were fed three times per week with freshly hatched Artemia nauplii. To avoid contamination with water-born bacteria, bacteria-free artificial seawater (FSW) was generated via vacuum filtration through a 0.2 μm Nylon membrane filter and used where noted.

### 2.2 Generation of Aposymbiotic Aiptasia

Aposymbiotic Aiptasia were generated at least three months prior to experimentation (Dani et al., 2016). Symbiotic Aiptasia were incubated in 0.008% menthol (from a 20% w/v menthol stock in 100% ethanol) in ASW for two weeks with daily menthol solution changes. After two weeks, aposymbiotic status was verified by lack of symbiont autofluorescence under fluorescence microscopy (Leica M165 FC), and aposymbiotic anemones were then maintained in ASW. Feeding was suspended during bleaching but resumed following confirmation of aposymbiotic state. During and after bleaching, aposymbiotic Aiptasia were maintained in darkness until at least three days prior to antibiotic treatment, at which time aposymbiotic Aiptasia were moved to the 12h:12h light:dark regimen.

### 2.3 Generation of Microbiome-depleted Aiptasia

Antibiotic solution (ABS) was prepared as described previously (Costa et al., 2021). Stock solutions of antibiotics consisted of the following: rifampicin (50 mg/ml in DMSO), nalidixic acid (50 mg/ml in MilliQ water), carbenicillin (100 mg/ml in MilliQ water), and chloramphenicol (50 mg/ml in 100% ethanol). ABS was used at a final concentration of 50 μg/ml of each antibiotic in ASW. The final ABS was vacuum filtered through a 0.2 μm Nylon membrane filter and stored in the dark at 4°C to be used within one month. Control and experimental Aiptasia were starved for two weeks prior to treatment to clear the digestive tract of microbiota associated with Artemia. Four ABS exposure conditions for each symbiotic state were used: 1) Aiptasia treated with 0.2 μm filter-sterilized ASW (FSW) for two weeks (control, C), 2) Aiptasia treated with ABS for seven days then washed twice in FSW (R0), 3) Aiptasia treated with ABS for seven days followed by two days of recovery in FSW (R2), and 4) Aiptasia treated with ABS for seven days followed by seven days of recovery in FSW (R7). ABS and FSW were changed daily over the course of the experiment.

### 2.4 Quantification of Colony-forming Units

To quantify the bacterial load of control and antibiotic-treated Aiptasia, Colony Forming Units (CFUs) were quantified for eight individuals per symbiotic state per treatment (n=32). Briefly, individuals were placed in 100 μl FSW in individual microcentrifuge tubes, lysed with a sterile pestle, and serially diluted to 1:1000 (i.e., 1:1, 1:10, 1:100, 1:1000). 1 μl of each dilution was plated on Marine Agar. Eight negative controls of 1 μl of FSW were additionally plated. All plates were incubated at 25°C for 48-72h, and colonies were counted to calculate total CFUs for each individual (accounting for dilution factor and total volume of lysis). To normalize for bacteria present in the FSW, CFUs counted on the negative control FSW-alone plates were averaged, and this average was multiplied by 100 to account for the total volume of FSW used in the anemone lysis. This number was subtracted from the total CFUs calculated for each individual. CFUs across treatments within symbiotic states were compared using Kruskal-Wallis and Dunn’s post-hoc tests in the FSA package v.0.10.0 (Ogle et al., 2025).

### 2.5 Quantification of Symbiosis by Red Channel Intensity

To determine changes in pigmentation (a proxy for symbiont density) over the course of antibiotic treatment in symbiotic Aiptasia, red channel intensity was measured for fifteen individuals from each condition at the beginning and end of the experiment. Photographs of each anemone (taken in standardized conditions with an iPhone 15 Pro at 2x magnification) were white-balanced in Adobe Photoshop (v.26.5.0). Images were then analysed in Matlab (v.R2025a) following the Coral Colour Intensity Analysis Utility 1.0 program (Winters et al., 2009) to extract red values (R-values). Three points from anemone tentacles were randomly chosen, measured, and averaged for each image. The relative change in R-value per anemone (measured as (final-initial)/initial) was compared across treatments using an analysis of variance and Tukey’s post-hoc test for pairwise comparisons (R Core Team, 2023).

### 2.6 Generation of 16S Gene Sequencing Microbiome Libraries

To characterize bacterial communities associated with microbiome depletion and recovery in symbiotic and aposymbiotic Aiptasia, 5-7 individuals from each treatment condition (C, R0, R2, R7) were prepared for 16S gene sequencing (n=48). Metabarcoding libraries were generated using a series of PCR amplifications for the V4/V5 region of the bacterial 16S rRNA gene (Apprill et al., 2015; Parada et al., 2016) as follows: 95°C for 40 sec, 58°C for 120 sec, and 72°C for 60 sec for 33 cycles, with a final elongation step of 72°C for 5 min. PCR products were purified using GeneJET PCR Purification kits (ThermoFisher) and eluted in 30 µl. Each PCR product was barcoded via five PCR cycles and visualized on a 1% agarose gel to assess relative concentrations. Five negative controls using sterile water were prepared and later used to remove contaminating sequences. Samples were pooled in equal concentrations, gel-extracted, and submitted for paired-end 250 bp sequencing on an Illumina Miseq (i100) at Tufts University Core Facility.

### 2.7 Analysis of Bacterial Community

All analyses for this section were performed in RStudio (v.4.4.0) (Posit team, 2025; R Core Team, 2023) following previously published work (Da-Anoy et al., 2025; Valadez-Ingersoll et al., 2025). 16S primers were removed from raw reads using cutadapt (Martin, 2011). DADA2 v.1.24.0 (Callahan et al., 2016) was used to perform quality filtering (including removing reads of off-target length and chimeras), resulting in the identification of 1,419 amplicon sequence variants (ASVs). Taxonomy was assigned at 100% sequence identity using the Silva v.138.1 database (Quast et al., 2013). 83 ASVs matching mitochondria, chloroplasts, or non-bacterial kingdoms were removed. Finally, decontam v.1.26.0 (Davis et al., 2018) was used to remove 87 contaminating ASVs based on co-occurrence in the negative controls. Underrepresented ASVs (ASVs in less than 0.02% of counts) were removed using MCMC.OTU v.1.0.10 (Matz, 2016), resulting in a total of 264 ASVs across samples for use in downstream analyses.

The effect of treatment on bacterial community composition, separated by symbiotic state, was assessed via Principal Coordinate Analyses (PCoA), and a PERMANOVA was used to compare Aitchison distances between samples by modeling the combined effect of treatment and symbiotic state (Martinez Arbizu, 2017; Oksanen et al., 2024).

Using vegan v.2.7-0 (Oksanen et al., 2024), cleaned counts were rarefied to 5,617 counts per sample. Alpha diversity metrics (Inverse Simpson’s Index, Shannon Index, ASV Richness, and Evenness) were used to further assess bacterial communities across treatments using phyloseq v.1.50.0 (function *estimate_richness* (McMurdie & Holmes, 2013)). The Inverse Simpson’s Index values were natural-log-normalized. The interacting effects of treatment and symbiotic state were measured using a two-way ANOVA for each Alpha diversity metric. Tukey’s post hoc tests were then used for pairwise comparisons (R Core Team, 2023).

To explore the relative abundance of different taxa across treatment groups and symbiotic states, the rarefied data were subsetted to contain ASVs within the Phylum Proteobacteria, and the relative abundance of each ASV was calculated within each sample and within each treatment group (for each symbiotic state). Proteobacteria relative abundances across treatment and symbiotic state – separated by Family – were visualized in a bar plot (McMurdie & Holmes, 2013). Counts from all ASVs in the Family Endozoicomonadaceae were summed for each sample and the relative abundance of Endozoicomonadaceae ASVs was plotted to compare across treatment conditions and symbiotic states. Summary statistics were computed (Wickham et al., 2023) for each condition and compared using a two-way ANOVA and Tukey post-hoc test (R Core Team, 2023).

Finally, DESeq2 v.1.44.0 (Love et al., 2014) was used to identify ASVs with differential abundance across treatments and symbiotic states. Pairwise comparisons were performed between each treatment condition and the control from the respective symbiotic state, and ASVs with significantly different abundances were identified with a *p*-adjusted value cutoff of 0.05. The log_2_-transformed abundance of each differentially abundant ASV in each anemone relative to the mean ASV abundance across all anemones was visualized in a heatmap (Kolde, 2019), and Phylum was noted.

### 2.8 RNA Extraction

To examine host gene expression, we performed TagSeq on symbiotic and aposymbiotic Aiptasia from each antibiotic treatment condition (as in Valadez-Ingersoll et al., 2024). Four to six individuals of each treatment/symbiotic state (n=40 total anemones) were flash frozen. Total RNA was extracted from each anemone by grinding whole anemones with sterile pestles during tissue lysis, followed by centrifugation at 13,000 rpm for 15 min at 4°C. The supernatant for each sample was processed using the RNAqueous Total RNA Isolation Kit (Invitrogen) according to the manufacturer’s instructions. RNA quantity and integrity were assessed using a DeNovix DS11+ Spectrophotometer. Samples were normalized to 10 ng/μl and submitted to the University of Texas at Austin Genomic Sequencing and Analysis Facility for TagSeq Library Preparation and sequencing on a NextSeq 1000.

### 2.9 Analysis of RNA-sequencing Data

Raw fastq reads were processed following an established pipeline (https://github.com/z0on/tag-based_RNAseq). Briefly, fastx_toolkit trimmed adapters and poly(A)+ tails, PCR duplicates were removed, and short (< 20 bp) and low-quality reads (quality score of ≤20) were discarded (Hannon, 2010). Cleaned reads were mapped to concatenated *E. pallida* (Baumgarten et al., 2015) and *S. linucheae* (González-Pech et al., 2021) transcriptomes using Bowtie2 (Langmead & Salzberg, 2012). A raw counts file containing all genes mapping exclusively to Aiptasia was used in downstream analysis (Supplementary Dataset 1). Genes mapping to the symbiont *S. linucheae* reference transcriptome were not analysed due to low counts.

All downstream analyses of RNA-sequencing data were performed in R v.4.4.0 (Posit team, 2025; R Core Team, 2023). DESeq2 v.1.44.0 (Love et al., 2014) identified differentially expressed genes (DEGs) between pairwise comparisons of interest by modelling all samples by condition (combined factor of symbiotic state and antibiotic treatment). ArrayQualityMetrics (v.3.62.0) (Kauffmann et al., 2009) tested for outliers, identifying one aposymbiotic R2 individual as an outlier, which was subsequently removed. Individual contrasts between treatments within each symbiotic state and between symbiotic states within each treatment identified DEGs (FDR-adjusted-*p*-value < 0.05) for 16 pairwise comparisons. Count data were *rlog*-normalized, and overall gene expression patterns were compared across all conditions through a Principal Component Analysis (PCA) implemented in vegan (v.2.7-0) (Oksanen et al., 2024). Gene expression similarity was tested using PERMANOVA implemented with the pairwiseAdonis package v.0.4.1 (Martinez Arbizu, 2017). Differential enrichment of gene pathways following microbiome depletion and recovery within each symbiotic state was investigated by analysing Biological Process (BP) Gene Ontology (GO) term enrichment via the Mann-Whitney U tests (GO-MWU) on all genes in each of six pairwise comparisons (*aposymbiotic-R0:aposymbiotic-C; aposymbiotic-R2:aposymbiotic-C; aposymbiotic-R7:aposymbiotic-C; symbiotic-R0:symbiotic-C; symbiotic-R2:symbiotic-C; symbiotic-R7:symbiotic-C*) (Wright et al., 2015). GO terms that were significantly enriched (FDR-*p*-value < 0.1) were plotted in dendrograms. GO term delta rank values of BP GO terms enriched in at least two of the six comparisons were visualized in ggplot2 (v.3.5.1) (Wickham, 2016) to compare differences in gene expression pathways across treatments and symbiotic states.

To identify clusters of genes highly correlated to host condition (symbiotic state, normalized CFU, relative abundance of total Endozoicomonadaceae ASVs, and change in R-value (in symbiotic-only WGCNA)) and treatments (C, R0, R2, R7), a weighted gene co-expression network analysis (WGCNA) was used (Langfelder & Horvath, 2008). Because normalized CFU, relative abundance of total Endozoicomonadaceae ASVs, and change in R-value data were measured from different individual anemones, these data points were averaged across treatments and applied to the respective individuals that matched their treatment in the gene expression analysis. First, a WGCNA for all Aiptasia of both symbiotic states was performed using a signed network, a softPower of 7, a minimum module size of 95 genes, and a threshold of 0.25 to merge modules with similar eigengene expression. Trait data and treatments were correlated with module eigengene expression using Pearson Correlation Coefficients, and the Student asymptotic p-value was calculated for each pairwise correlation. Next, WGCNAs were performed separately for symbiotic and aposymbiotic Aiptasia following the same protocol (softPower=15, threshold=0.45) (Langfelder & Horvath, 2008; Radice et al., 2024). The grey module containing 12 unassigned genes was removed from the aposymbiotic-only WGCNA. For each WGCNA, pairwise correlations between traits/treatments and module eigengenes were visualized in heatmaps, and the numbers of genes included in each module was visualized in barplots (ggplot2 (v.3.5.1) (Wickham, 2016)). BP GO term enrichment analysis was performed on the genes included in each module using Fisher’s Exact Tests (gene presence/absence in the module) (Wright et al., 2015).

Significantly enriched GO terms (FDR-*p*-value < 0.1) were plotted in dendrograms, and modules enriched for immune terms were further explored (**Tables S1, S2,** and **S3**). In the symbiotic-only WGCNA, significantly enriched BP GO terms involved in immunity were identified in the Black module (**Table S2**). Genes annotated with these immune GO terms were subsetted from the Black module if they were identified as differentially expressed (FDR-*p*-value < 0.05 from DESeq2) in any treatment condition (R0, R2, or R7) compared to the control. In the aposymbiotic-only WGCNA, significantly enriched BP GO terms involved in immunity were identified in the Blue module (**Table S3**). Genes annotated with these immune GO terms were subsetted from the Blue module if they were identified as differentially expressed (FDR-*p*-value < 0.05 from DESeq2) in any treatment condition (R0, R2, or R7) compared to the control. Expression levels of these genes across samples were visualized using pheatmap v.1.0.12 (Kolde, 2019).

### 2.10 Western Blotting of NF-κB

Western blotting of NF-κB was performed as described previously on whole anemone extracts from three Aiptasia of each treatment and symbiotic state (Aguirre Carrión et al., 2023; Mansfield et al., 2017; Valadez-Ingersoll et al., 2024). Western blots were repeated three times with new samples for symbiotic Aiptasia and twice with new samples for aposymbiotic Aiptasia (total of 60 anemones). Briefly, individual anemones were homogenized in 60 μl of 2x SDS-sample buffer (0.125 M Tris-HCl [pH 6.8], 4.6% w/v SDS, 20% w/v glycerol, 10% v/v β-mercaptoethanol, 0.2% w/v bromophenol blue) using a pestle followed by heating at 95°C for 10 min. Samples were centrifuged for 10 min at 13,000 rpm and the supernatant was collected. Extracts were electrophoresed on 7.5% SDS-polyacrylamide gels, and proteins were transferred to nitrocellulose membranes. Membranes were blocked for 1 h in TBST (10 mM Tris-HCl [pH 7.4], 150 mM NaCl, 0.05% v/v Tween 20 containing 5% non-fat powdered milk), then incubated overnight at 4°C with primary rabbit anti-Ap-NF-κB antiserum (Mansfield et al., 2017) (diluted 1:5,000). Membranes were washed five times with TBST and incubated for 1 h with secondary goat-anti-rabbit-HRP conjugated antiserum (1:4,000; Cell Signaling Technology). After further washing, immunoreactive proteins were detected with SuperSignal West Dura Extended Duration Substrate (Fisher Scientific) and were imaged on a Sapphire Biomolecular Imager. Bands corresponding to Aiptasia NF-κB (Ap-NF-κB) were normalized to total protein loaded in each lane using Ponceau S stain and ImageJ. For each individual, normalized Ap-NF-κB values were compared to the average normalized control Ap-NF-κB for each respective blot. The relative values were plotted using ggplot (v.3.5.1) (Wickham, 2016), and the effects of antibiotic treatment and symbiotic state on relative normalized ap-NF-κB levels were measured using an analysis of variance (R Core Team, 2023).

## 3. RESULTS

### 3.1 Antibiotic Treatment Depletes Bacterial Load and Alters Pigmentation in Aiptasia

To measure the effect of antibiotic treatment on bacterial load in aposymbiotic and symbiotic Aiptasia, total colony forming units (CFUs) of bacteria isolated from anemones across treatment conditions and symbiotic states were quantified. Four treatment groups were used: 1) Control Aiptasia (C) incubated in FSW for two weeks; 2) Aiptasia treated with antibiotic solution (ABS) for seven days with no recovery period (R0); 3) Aiptasia treated with ABS for seven days followed by two days of recovery in FSW (R2); and 4) Aiptasia treated with ABS for seven days followed by seven days of recovery in FSW (R7). Directly after antibiotic treatment (R0), bacterial load was significantly decreased in symbiotic and aposymbiotic Aiptasia as compared to control (C) anemones (∼125-fold decrease in symbiotic and ∼1,500-fold decrease in aposymbiotic Aiptasia; FDR *p*-value of <0.05). In both symbiotic states, anemones were able to reestablish bacterial loads comparable to control anemones within two days (R2), and these levels were maintained for up to seven days after antibiotic treatment (R7) (**Fig. 1a**). Of note, the overall bacterial load was approximately 8-fold higher in aposymbiotic controls compared to symbiotic controls (FDR *p*-value <0.05).

**Figure 1.**
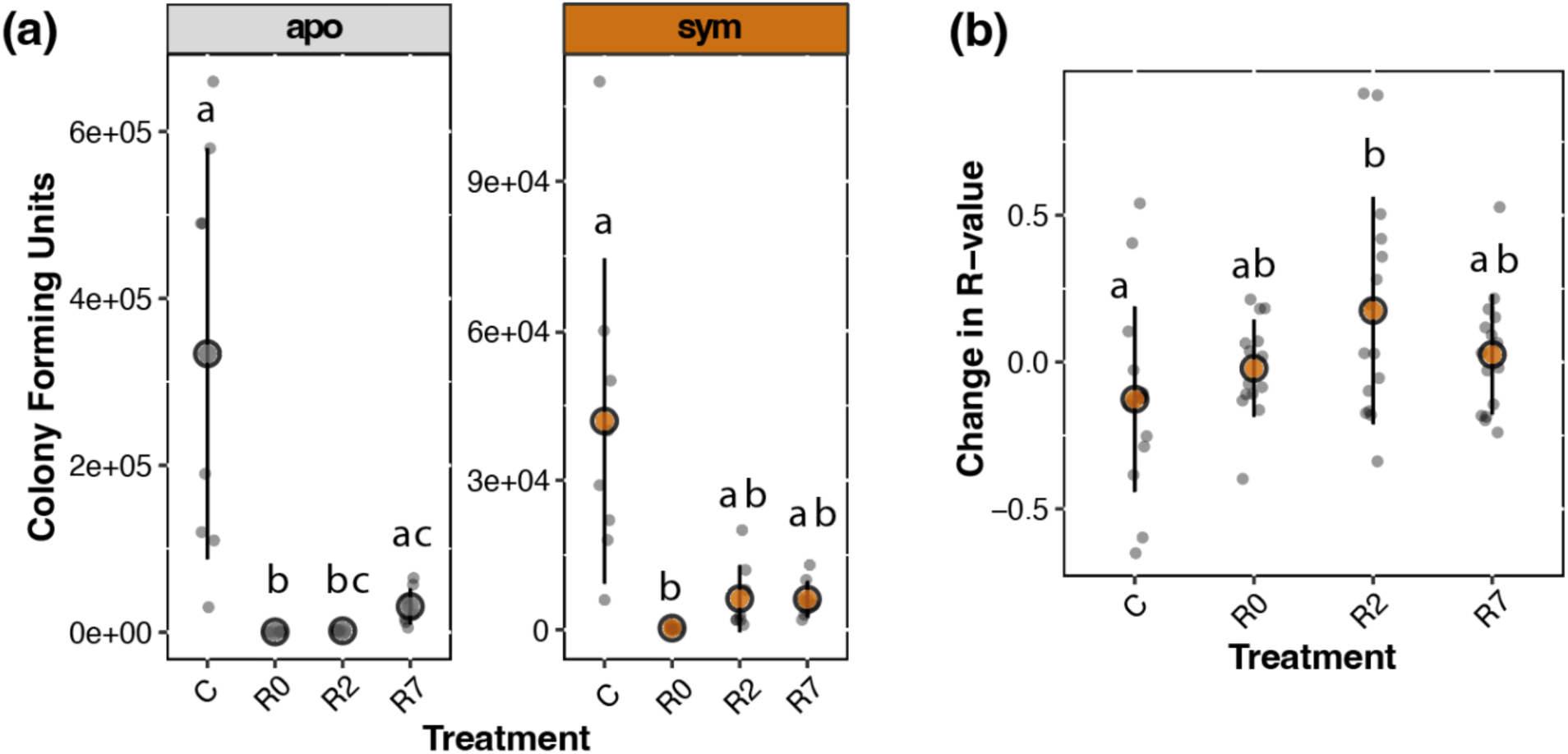
Antibiotic treatment results in depletion of bacterial load in symbiotic and aposymbiotic Aiptasia and reduced pigmentation in symbiotic anemones. **(a)** Total colony forming units (CFUs) from eight anemones per treatment per symbiotic state were quantified to determine bacterial load. In each symbiotic state, distinct letters represent significant differences between treatments (Kruskal-Wallis and Dunn’s post-hoc tests, *p*<0.05). **(b)** Relative change in red channel intensity (R-value, proxy for pigmentation) for 15 symbiotic individuals from each treatment as measured at the beginning and end of the experiment ((final-initial)/initial). Distinct letters represent significant differences between treatments (analysis of variance and Tukey’s post-hoc test, *p*<0.05), and higher values indicate reduction in pigmentation. In **(a)** and **(b)**, coloured circles (grey for apo and orange for sym) show the mean, and black bars indicate standard deviation.

To determine the impact of antibiotic treatment on algal symbiont densities in symbiotic Aiptasia, we measured red channel intensity (R-value, a proxy for symbiont density) of symbiotic Aiptasia at the beginning (i.e., before Day 1 of antibiotic addition for R0, R2, and R7, and before Day 1 of FSW for control anemones) and at the end (i.e., after rinsing in FSW on Day 7 of ABS for R0 anemones; after Day 2 of FSW for R2 anemones; after Day 7 of FSW for R7 anemones; after Day 14 of FSW for control anemones) of the experiment and calculated the change in R-value as (final-initial)/initial. A positive change in red channel intensity indicates a decrease in pigmentation and therefore a presumed loss of symbiont density (Winters et al., 2009). Based on the change in R-values, R2 anemones experienced some bleaching (mean change of 0.18 in R2 compared to -0.13 in control anemones; FDR *p*-value < 0.05), but anemones recovered from algal loss by R7 (the mean change of 0.026 in R7 was not significantly different from the mean change of -0.13 in control anemones) (**Fig. 1b**).

### 3.2 Microbial Depletion and Recovery Alter Alpha and Beta Diversity, but Differ by Symbiotic State

We profiled the bacterial communities of symbiotic and aposymbiotic Aiptasia from control (C), R0, R2, and R7 conditions (n = 48) (**Fig. S1**). A total of 2,250,769 sequences were acquired with a mean depth of coverage of 46,891 +/- 21,710 (mean +/- SD) per sample (**Table S4**). Sample rarefication yielded a total of 5,617 reads per sample with a total of 264 unique ASVs identified across samples.

To investigate how antibiotic treatment alters microbiome community composition following depletion and recovery, we compared microbiome community composition and calculated alpha diversity metrics for each treatment and symbiotic state. Principal Coordinate Analyses (PCoAs) tested differences in overall community composition across treatments and showed that community composition significantly differed by antibiotic treatment (*p_T_* < 0.001) in both symbiotic (**Fig. 2a**) and aposymbiotic (**Fig. 2b**) samples. Within both symbiotic states, all pairwise comparisons of Aitchison distances (a beta-dispersion index of sample dissimilarity) among antibiotic treatments were significant (FDR *p*-value < 0.05) (**Fig. 2a** and **2b**).

**Figure 2.**
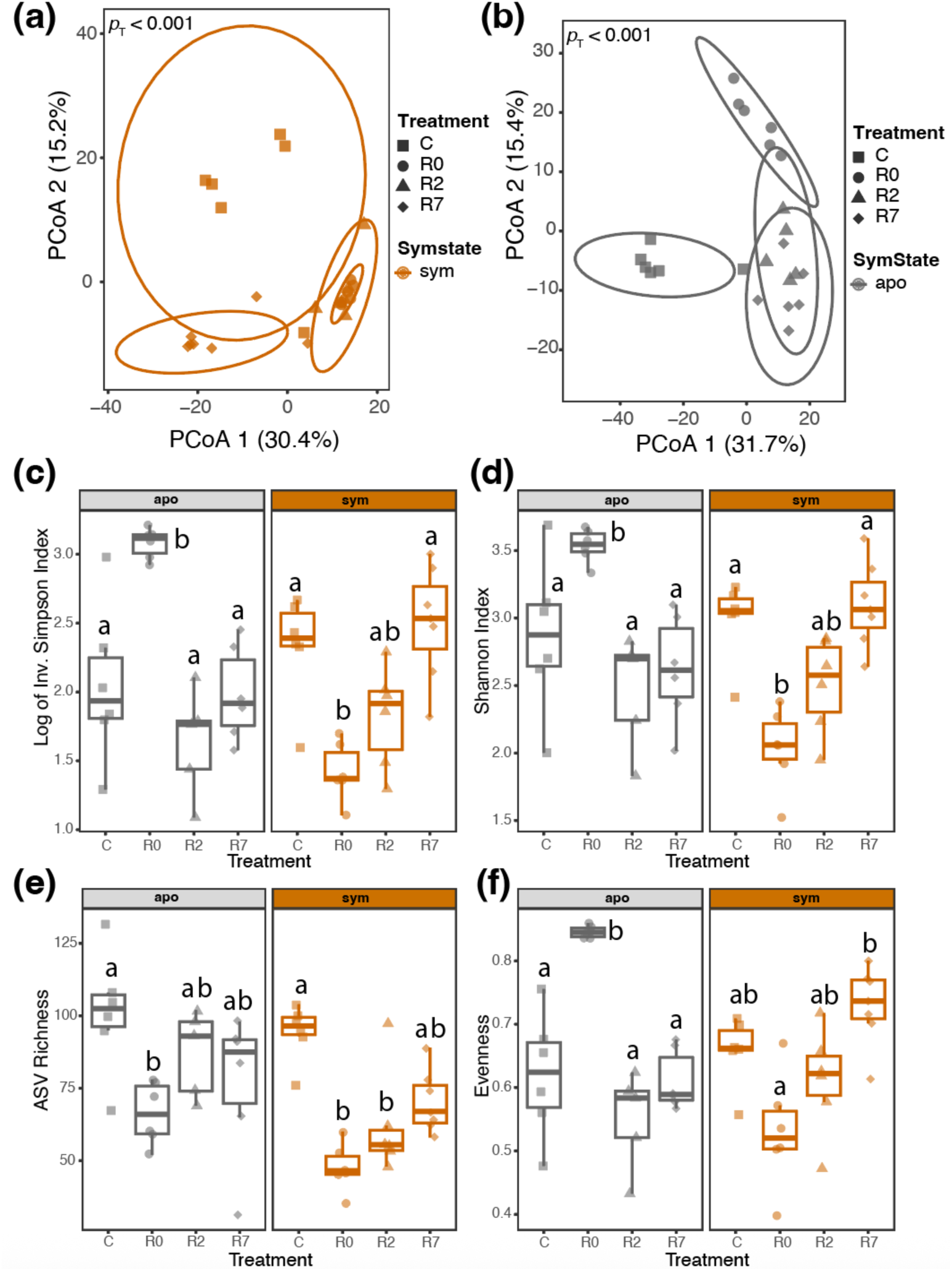
Microbiome depletion and recovery alter diversity metrics in symbiotic and aposymbiotic Aiptasia. PCoAs display the similarity in bacterial communities from all ASVs in 16S data from symbiotic **(a)** and aposymbiotic **(b)** Aiptasia across antibiotic treatments (shapes). PERMANOVA results for the effect of treatment (*p*T) for each symbiotic state are included in **(a)** and **(b)**. Alpha diversity metrics (Natural log of the Inverse Simpson’s Index **(c)**; Shannon Index **(d)**; Richness **(e)**; and Evenness **(f)**) of rarefied 16S data were measured across symbiotic states and treatment conditions. In each symbiotic state, distinct letters represent significant differences in alpha diversity metrics between treatments (ANOVA and Tukey’s post-hoc tests, *p*<0.05). Box plots show the median and the 1^st^ and 3^rd^ quartiles (hinges). Whiskers extend to the maximum and minimum points if they fall within 1.5x the Interquartile Range (if no whiskers are drawn, points outside of the box are outliers). Each point represents the alpha diversity metric of one anemone, with shapes representing antibiotic treatments and colour representing symbiotic state, as in **(a)** and **(b)**.

Alpha diversity between antibiotic treatments within each symbiotic state was measured by calculating the natural log of the Inverse Simpson’s Index, the Shannon Index, Richness, and Evenness. The Inverse Simpson’s Index and the Shannon Index account for ASV richness (number of ASVs within each sample) and evenness (how evenly distributed the ASVs are within each sample), with higher values indicating greater diversity. Treatment (*p* < 0.05) and the interaction between treatment and symbiotic state (*p* < 0.001) both had significant effects on Simpson’s Index (**Fig. 2c**) and Shannon Index (**Fig. 2d**) values. Within aposymbiotic Aiptasia, R0 anemones had higher Simpson’s and higher Shannon Indices as compared to all other treatments (*p* < 0.05; **Fig. 2c** and **Fig. 2d**). Within symbiotic Aiptasia, R0 anemones had lower Simpson’s and lower Shannon Indices than control and R7 (*p* < 0.05), but not R2 samples (**Fig. 2c** and **Fig. 2d**). Treatment (*p* < 0.001) and symbiotic state (*p* < 0.01), but not their interaction, had significant effects on ASV Richness (**Fig. 2e**). Within aposymbiotic Aiptasia, Richness was lower in R0 compared to the control (*p* < 0.05; **Fig. 2e**). Within symbiotic Aiptasia, Richness was lower in R0 and R2 compared to the control (*p* < 0.05; **Fig. 2e**). Finally, treatment (*p* < 0.01) and the interaction between treatment and symbiotic state (*p* < 0.001) had significant effects on Evenness (**Fig. 2f**). Within aposymbiotic Aiptasia, Evenness was highest in R0 but returned to control levels by R2 (*p* < 0.05; **Fig. 2f**). Within symbiotic Aiptasia, Evenness was higher in R7 samples than in R0 samples, but no other comparisons were significant (*p* < 0.05; **Fig. 2f**). Overall, the alpha diversity results indicate that the dynamics of microbiome reestablishment following antibiotic treatment differ by symbiotic state.

### 3.3 Relative Abundance and ASV Counts of Endozoicomonadaceae Decrease in Aposymbiotic but Increase in Symbiotic Aiptasia Following Antibiotic Treatment

To investigate how specific microbial taxa associated with symbiotic and aposymbiotic Aiptasia were altered following antibiotic treatment and recovery, we compared the relative abundance of different taxa within the Phylum Proteobacteria. In aposymbiotic Aiptasia, the relative abundance of the Family Endozoicomonadaceae decreased following antibiotic treatment (from ∼38% of the total ASVs in control anemones to ∼0.03% in R0 anemones) and remained low even in R7 samples (∼0.18% of the total ASVs), suggesting that their relative abundances failed to recover (**Fig. 3a** and **3b**; *p* < 0.001). However, the relative abundance of the Family Marinobacteraceae increased throughout the course of microbiome reestablishment in aposymbiotic anemones (R2 and R7; **Fig. 3a**). Interestingly, in symbiotic anemones, the relative abundance of Endozoicomonadaceae increased following antibiotic treatment at R0 and R2 (25% in control compared to ∼76% in R0 and ∼60% in R2) (**Fig. 3a** and **3b**; *p* < 0.01). While the relative abundance of Endozoicomonadaceae was lower in symbiotic R7 anemones compared to symbiotic control anemones (25% in control compared to ∼2.7% in R7), this difference was not significant (**Fig. 3a** and **3b**).

**Figure 3.**
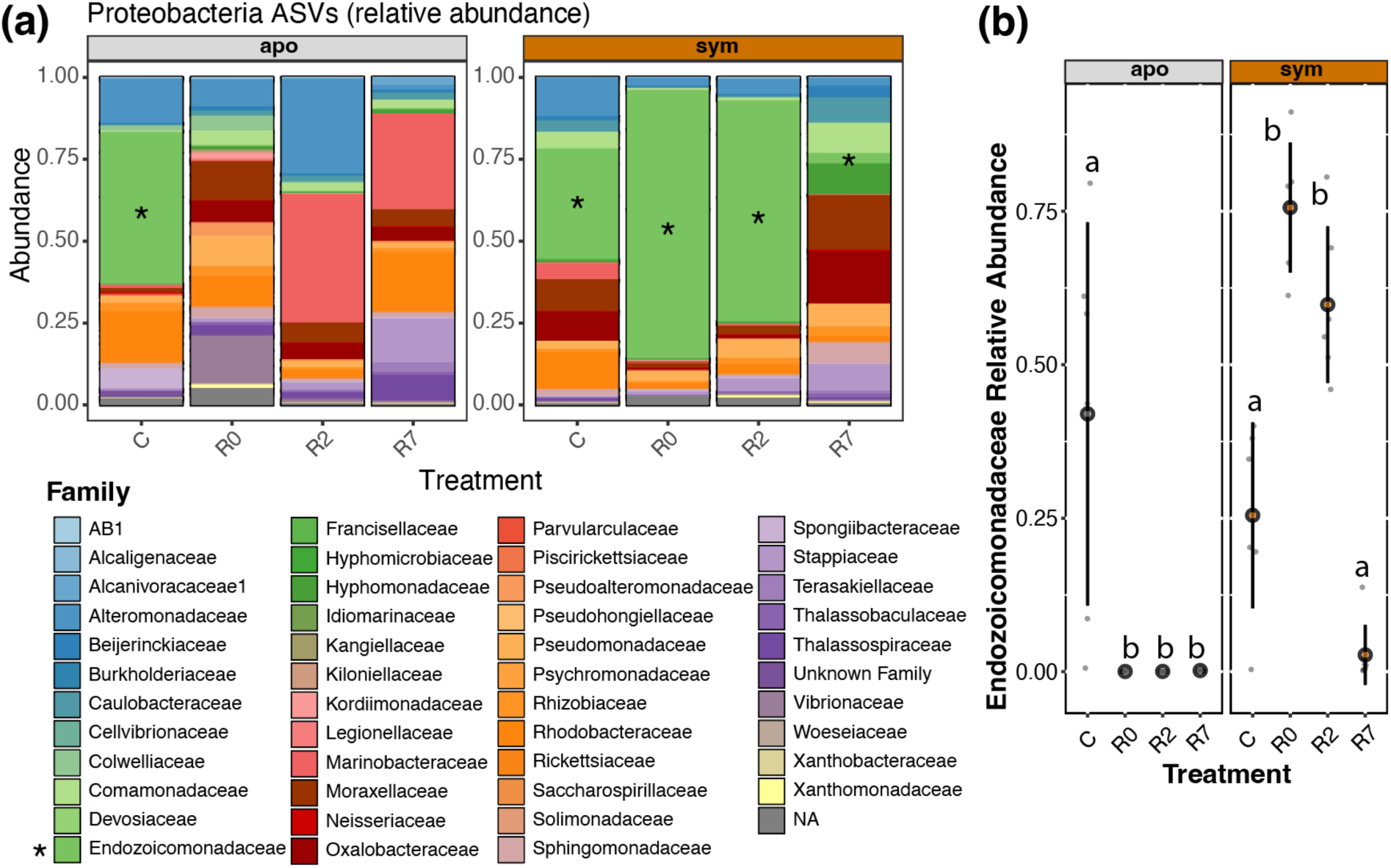
Relative abundances of bacterial Families are impacted by antibiotic treatment and recovery. **(a)** The relative abundance of 16S ASVs within the Phylum Proteobacteria was measured across antibiotic treatments and symbiotic state (apo=aposymbiotic; sym=symbiotic). Colours in the stacked bar graph correspond to different bacterial Families. * denotes Endozoicomonadaceae in the legend and in the bar graphs (absent from aposymbiotic R0, R2, and R7 because the relative abundance is <1%. **(b)** The relative abundance of the four ASVs in the Family Endozoicomonadaceae compared to all 264 ASVs was compared across antibiotic treatments and symbiotic states. Coloured circles (grey for apo and orange for sym) show the mean and black bars indicate standard deviation. Within each symbiotic state, distinct letters represent significant differences in Endozoicomonadaceae relative abundance across treatments (ANOVA and Tukey’s post-hoc tests (*p*<0.01)).

### 3.4 ASV Composition Recovers in Symbiotic but not Aposymbiotic Aiptasia

To determine how individual ASVs were impacted by antibiotic treatment and recovery, we performed differential abundance analyses using DESeq2 (Love et al., 2014). Within each symbiotic state, we performed pairwise comparisons between each treatment and the control. In symbiotic samples, there were 47 differentially abundant ASVs between R0 and control samples, 40 between R2 and control samples, and 16 between R7 and control samples (FDR *p*-value of < 0.05 for each pairwise comparison), suggesting that bacterial communities return to baseline during recovery (**Fig. 4**). In aposymbiotic samples, there were 16 differentially abundant ASVs between R0 and control samples, 8 between R2 and control samples, and 25 between R7 and control samples (FDR *p*-value of <0.05 for each pairwise comparison). Although there were fewer overall differentially abundant ASVs in aposymbiotic anemones than in symbiotic anemones, After the seven-day recovery period, community composition in apopsymbiotic anemones was more distinct from control communities than it was directly following antibiotic treatment (R0), indicating that community composition did not begin to return to baseline composition as quickly as it did in symbiotic Aiptasia (**Fig. 4**).

**Figure 4.**
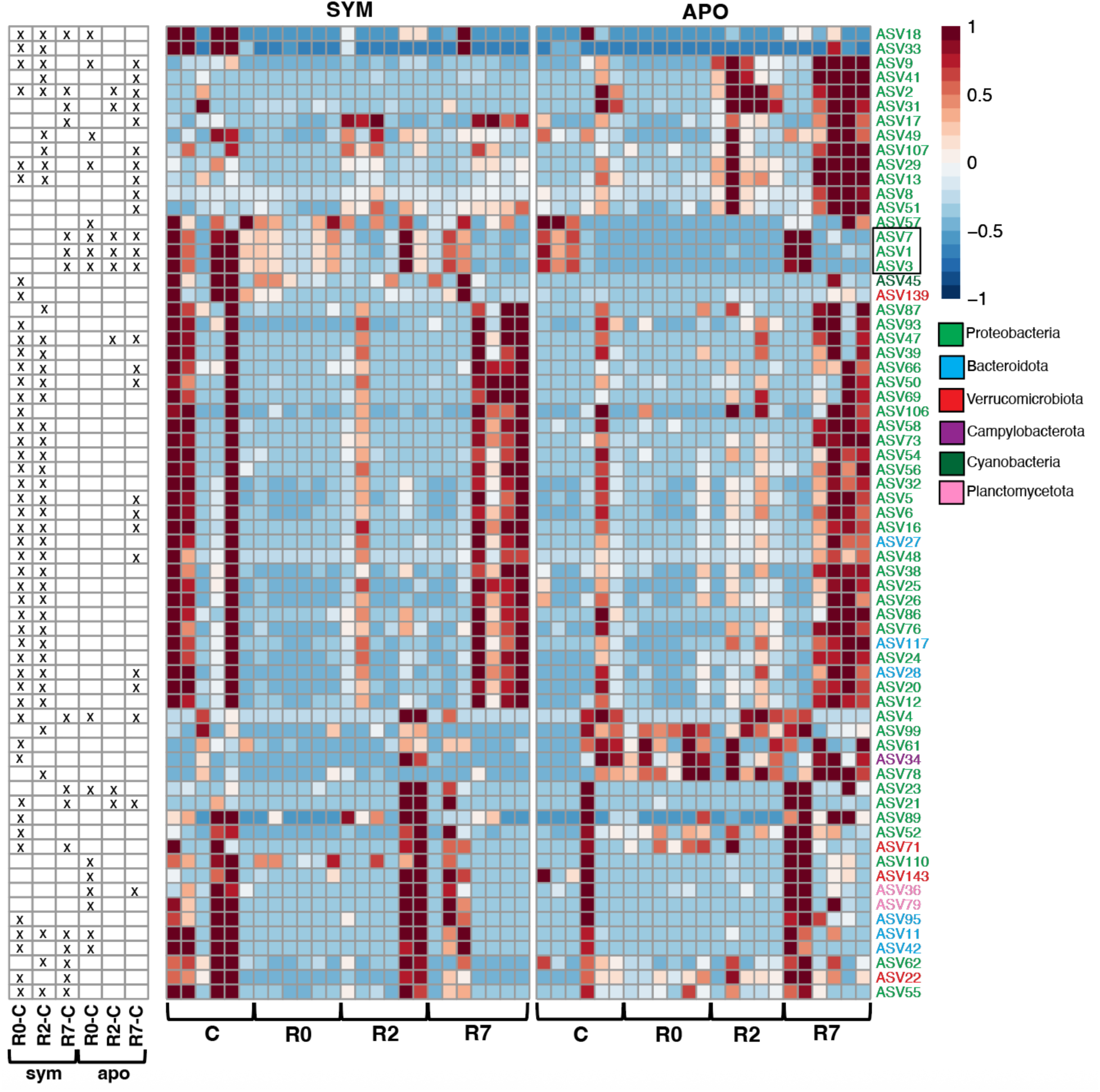
Similar bacterial communities reestablish in symbiotic, but not aposymbiotic, Aiptasia following antibiotic treatment. ASVs with differential abundance in the indicated treatment conditions compared to the control (within the respective symbiotic state) were plotted in a heatmap with the colour scale representing the log_2_-transformed abundance of each ASV (row) in each anemone (column) relative to the mean ASV abundance across all anemones. ASVs are hierarchically clustered (dendrogram not shown) based on similar abundance across samples and are coloured by their Phylum. A box is drawn around ASVs in the Family Endozoicomonadaceae. In the left panel, **x** indicates which ASVs have significantly different abundances in each treatment compared to the respective control (FDR *p*-value<0.05 from DESeq2).

Differentially abundant ASVs across comparisons were dominated by Proteobacteria. In agreement with **Fig. 3a** and **3b**, three of the four Endozoicomonadaceae ASVs (ASVs 1, 3, and 7) had lower abundances in R0, R2, and R7 samples compared to control samples in aposymbiotic Aiptasia (FDR *p*-value < 0.05) (**Fig. 4**). In contrast to the results from **Fig. 3b**, individual Endozoicomonadaceae ASVs were not significantly higher in symbiotic R0 and R2 samples compared to control, but they were significantly lower in R7 samples (FDR *p*-value < 0.05) (**Fig. 4**). The decrease in Endozoicomonadaceae immediately following antibiotic treatment in aposymbiotic but not symbiotic Aiptasia suggests that Endozoicomonadaceae in symbiotic anemones are buffered from antibiotic-induced loss because of an association with the algal symbionts.

### 3.5 Immune System Pathways are Downregulated During Microbiome Reestablishment in both Symbiotic and Aposymbiotic Aiptasia

To investigate the influence of microbiome depletion and recovery on Aiptasia gene expression, overall gene expression patterns across antibiotic treatments and symbiotic states were compared using a Principal Component Analysis (PCA). The PCA revealed a significant effect of symbiotic state on gene expression, displayed by separation of samples along PC1 (*p_Sym_* < 0.001), which explained 18.5% of the variation (**Fig. 5a**). Antibiotic treatment also had a significant effect on gene expression, revealed by separation of samples along PC2 (*p_T_* < 0.001), which explained 7.7% of the variation (**Fig. 5a**). Additionally, the interaction between treatment and symbiotic state was significant (*_pT*Sym_* < 0.001).

**Figure 5.**
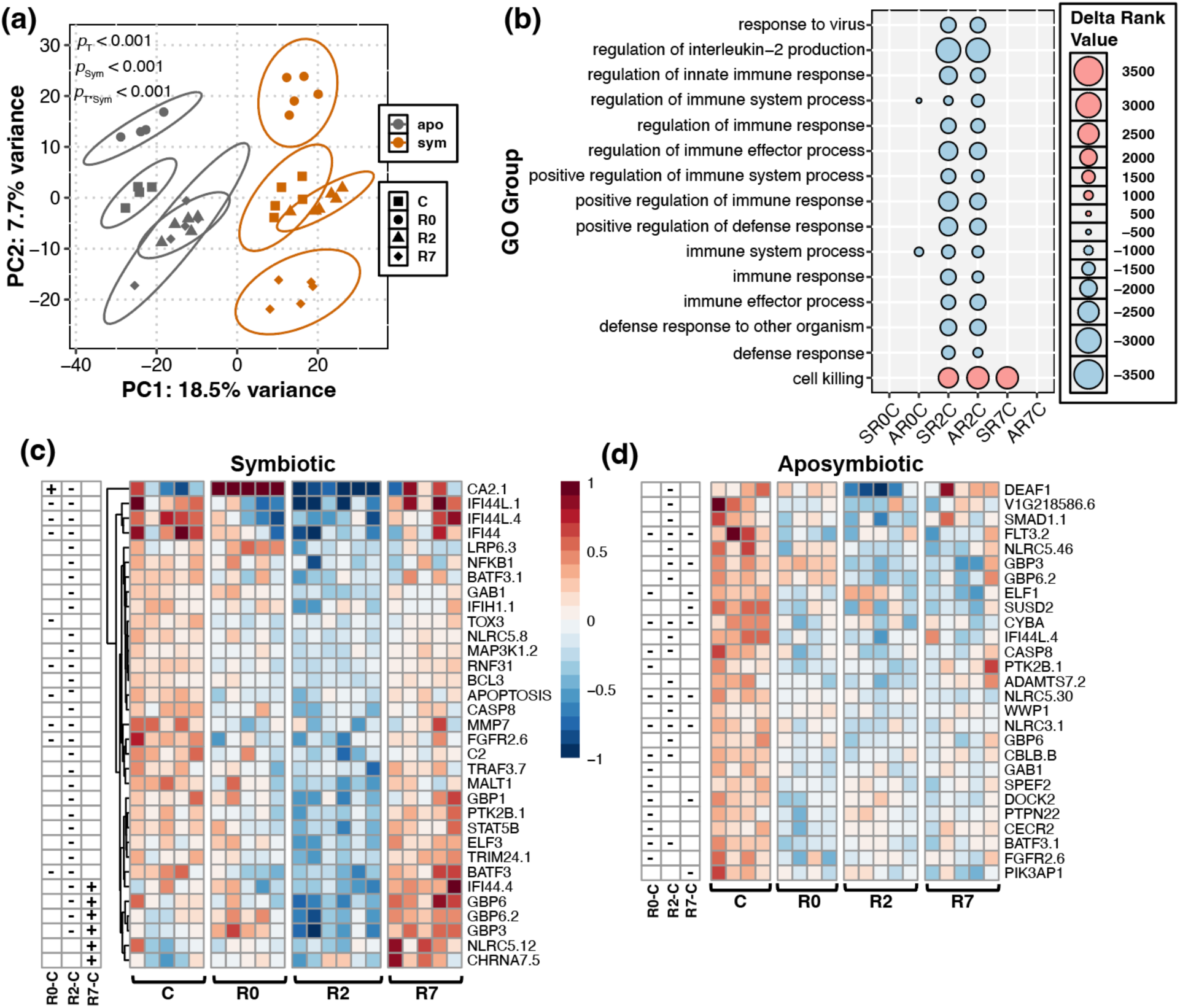
Microbiome depletion and reestablishment modifies host immune gene expression in symbiotic and aposymbiotic Aiptasia. **(a)** Principal component analysis (PCA) of log2**-** transformed counts for symbiotic (orange) and aposymbiotic (grey) Aiptasia across antibiotic treatments (shapes). PERMANOVA results for the effect of antibiotic treatment (*p_T_*), symbiotic state (*p_Sym_*) and the interaction between treatment and symbiotic state (*_pT*Sym_*) are included. **(b)** Delta rank values (represented by point size) of enriched Gene Ontology terms related to immunity were compared across each treatment condition relative to the control (within symbiotic state). Terms were plotted if they were significantly enriched in the treatment (red, positive Delta Rank Values) or enriched in the control (blue, negative Delta Rank Values) in at least two comparisons. SR0C=symbiotic R0 to symbiotic control; AR0C=aposymbiotic R0 to aposymbiotic control; SR2C=symbiotic R2 to symbiotic control; AR2C=aposymbiotic R2 to aposymbiotic control; SR7C=symbiotic R7 to symbiotic control; AR7C=aposymbiotic R7 to aposymbiotic control. **(c)** In symbiotic Aiptasia, genes were identified within the Black WGCNA module that were annotated with significantly enriched immune GO terms and that were differentially expressed between R0 and control, R2 and control, or R7 and control. **(d)** In aposymbiotic Aiptasia, genes were identified within the Blue WGCNA module that were annotated with significantly enriched immune GO terms and that were differentially expressed between R0 and control, R2 and control, or R7 and control. In **(c)** and **(d)**, the left panels indicate which genes are differentially expressed (**+** represents positive differential expression in the treatment compared to the control; **-** represents negative differential expression in the treatment compared to the control; FDR *p*-value <0.05 from DESeq2).

To identify differentially expressed genes (DEGs) between the control and antibiotic treatments in each symbiotic state, we performed pairwise comparisons of gene expression. In symbiotic Aiptasia, there were 1,000 DEGs between R0 and control samples, 983 DEGs between R2 and control samples, and 1,347 DEGs between R7 and control samples (FDR *p*-value < 0.05). In aposymbiotic Aiptasia, there were 342 DEGs between R0 and control samples, 958 DEGs between R2 and control samples, and 814 DEGs between R7 and control samples (FDR *p*-value < 0.05). We additionally determined the number of DEGs for all other possible pairwise comparisons within and between symbiotic states (**Table S5**).

To explore biological responses related to microbiome depletion and recovery across symbiotic states, we performed Gene Ontology (GO) enrichment analysis on terms in the Biological Process (BP) GO category. GO terms enriched between each antibiotic treatment and the control within each symbiotic state were identified. Notably, symbiotic and aposymbiotic R2 samples, and aposymbiotic (and to a lesser degree symbiotic) R0 samples were positively enriched for functions relating to metabolism (e.g., “cellular lipid metabolic process;” “cofactor metabolic process;” “monosaccharide metabolic process) (FDR *p-*value < 0.1; **Table S6**). For both symbiotic states, 14 terms related to immune function (e.g., “regulation of interleukin-2 production;” “immune response”) were decreased in R2 compared to control samples, indicating that two days following antibiotic treatment, Aiptasia had decreased gene expression pathways related to immunity regardless of symbiotic state (**Fig. 5b**).

We also used WGCNAs to investigate the correlation between gene expression and experimental treatments/host traits following microbiome treatment and recovery. In the combined symbiotic and aposymbiotic WGCNA, 16,345 genes were assigned to 10 modules (**Fig. S2**). The Blue module (containing 4,348 genes) was enriched for GO terms related to immunity (**Table S1**). This module was positively correlated with aposymbiotic anemones, control, R0, and R7 treatments in aposymbiotic Aiptasia, as well as normalized CFU counts (**Fig. S2**). This module was negatively correlated with symbiotic anemones, the R2 treatment in symbiotic Aiptasia, the R2 treatment in general, and relative abundance of Endozoicomonadaceae ASVs (**Fig. S2**).

Because of the significant effect of symbiotic state on overall patterns of gene expression, transcriptomic signals specific to antibiotic treatment may be masked in the combined WGCNA model. To account for this potential masking, we performed separate WGCNAs on symbiotic and aposymbiotic samples. In the symbiotic-only WGCNA, 16,283 genes were assigned to 16 modules (**Fig. S3**). The Black module (containing 1,659 genes) was enriched for GO terms related to immunity, including the NF-κB pathway (**Table S2**). This module was positively correlated with the R7 treatment in symbiotic Aiptasia (**Fig. S3**) and negatively correlated with the R2 treatment, relative abundance of Endozoicomonadaceae ASVs, and change in R-value in symbiotic Aiptasia (**Fig. S3**). 941 Black module genes were annotated with immune GO terms, 10 of which were differentially expressed in symbiotic R0 compared to symbiotic control Aiptasia (1 upregulated; 9 downregulated). 30 immune genes in the Black module were differentially expressed in symbiotic R2 compared to symbiotic control Aiptasia (all downregulated), and 6 immune genes in the Black module were differentially expressed in symbiotic R7 compared to symbiotic control Aiptasia, all of which were upregulated (**Fig. 5c**).

In the aposymbiotic-only WGCNA, 16,920 genes were assigned to 13 modules (**Fig. S4**). The Blue module (containing 3,429 genes) was enriched for GO terms related to immunity. While terms relating explicitly to the NF-κB pathway were not identified, the term “Response to Tumor Necrosis Factor” was identified (**Table S3**). This module was positively correlated with the control treatment, normalized CFU counts, and the relative abundance of Endozoicomonadaceae ASVs in aposymbiotic Aiptasia (**Fig. S4**). 565 Blue module genes annotated with immune GO terms were identified, 15 of which were differentially expressed in aposymbiotic R0 compared to aposymbiotic control Aiptasia (all downregulated). 17 immune genes in the Blue module were differentially expressed in aposymbiotic R2 compared to aposymbiotic control Aiptasia (all downregulated), and 9 Blue module genes were differentially expressed in the aposymbiotic R7 compared to the aposymbiotic control Aiptasia, all of which were downregulated (**Fig. 5d**).

Symbiotic Aiptasia had a stronger transcriptomic response than aposymbiotic Aiptasia, including in expression of immune genes. However, both symbiotic states differentially regulated similar immune system pathways during microbiome reestablishment. Immune pathway genes downregulated in R2 anemones included genes involved in the NF-κB pathway (e.g., *NFKB1, BCL3, IFI44.4*, and *ELF3* in symbiotic Aiptasia) (Cleves et al., 2020; Valadez-Ingersoll et al., 2024), as well as other immune pathway genes downregulated throughout the course of recovery. Other potential immune system pathways involved in microbiome reestablishment in both symbiotic states include C-Type Lectin Receptor signaling (*MALT1, BCL3*), Tumor Necrosis Factor signaling (*TRAF3.7, BCL3, CYBA, CASP8, PTK2B.1, ADAMTS7.2*), NOD-Like Receptor signaling (*TRAF3.7, RNF31*), and the MAPK pathway (*MAP3K1.2, FGFR2.6*) (Jacobovitz et al., 2021). Many of these immune pathway genes were no longer downregulated following seven days of recovery (R7), especially in symbiotic Aiptasia (**Fig. 5c** and **5d**).

**3.6 Protein Levels of the Active Form of NF-κB Increase in Microbially Depleted Aiptasia** Because gene expression pathways related to immunity were downregulated following microbiome depletion in Aiptasia, we directly tested the effect of microbiome depletion on protein levels of NF-κB – a transcription factor associated with immunity and symbiosis with intracellular algae. In both symbiotic and aposymbiotic Aiptasia, the levels of the processed (active) form of the NF-κB protein were higher in R0, R2, and R7 samples compared to the levels in the control anemones of the respective symbiotic state (**Fig. 6a** and **6b**). In symbiotic Aiptasia, processed NF-κB levels were ∼2.3 times higher in R0 samples compared to the control, ∼2.3 times higher in R2 samples compared to the control, and ∼1.8 times higher in R7 samples compared to the control (**Fig. 6a** and **6c**). In aposymbiotic Aiptasia, processed NF-κB levels were ∼2.7 times higher in R0 samples compared to the control, ∼4.6 times higher in R2 samples compared to the control, and ∼2.7 times higher in R7 samples compared to the control (**Fig. 6b**). Therefore, antibiotic treatment had a significant effect on levels of processed NF-κB regardless of symbiotic state (*p*<0.05; **Fig. 6c**) These results indicate that, while general immunity gene expression is downregulated following antibiotic treatment, levels of transcription factor NF-κB are increased.

**Figure 6.**
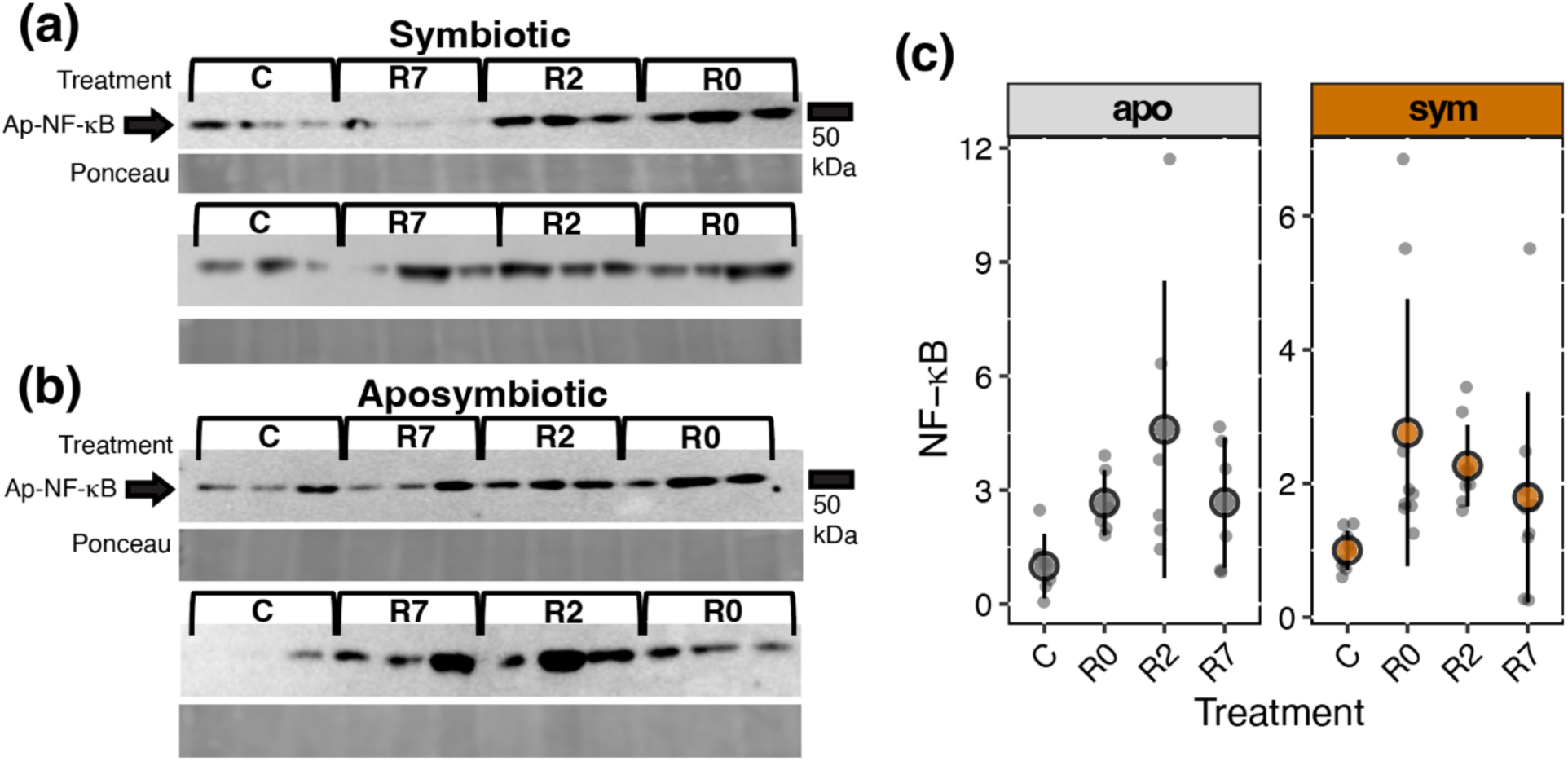
NF-κB protein levels are increased in symbiotic and aposymbiotic Aiptasia following microbiome depletion. Western blots of NF-κB in symbiotic **(a)** and aposymbiotic **(b)** Aiptasia across antibiotic conditions. Two representative blots for each symbiotic state are shown. Three independent anemones were analysed for each treatment in each Western blot. NF-κB levels were normalized to Ponceau staining for each sample. Location of the 50 kDa molecular weight marker is indicated to the right of the blots. **(c)** Levels of the processed form of NF-κB were normalized to total protein (Ponceau) for each anemone, and this value was normalized to the average control NF-κB:Ponceau value for each respective blot then plotted. Coloured circles (grey for apo and orange for sym) show the mean and black bars indicate standard deviation.

## 4. Discussion

We investigated the tripartite interactions between cnidarians, their intracellular photosynthetic algae, and their bacterial microbiome using the facultatively symbiotic sea anemone Aiptasia with the overall goal of better understanding how the cnidarian holobiont responds to environmental stress. Using an antibiotic cocktail, we depleted the microbiomes of symbiotic and aposymbiotic Aiptasia and investigated how the microbiome reestablishes using 16S microbiome characterization and gene expression profiling of the anemone host. Shifts in alpha and beta diversity metrics indicated that microbiomes of aposymbiotic Aiptasia became more disrupted immediately following antibiotic treatment while symbiotic Aiptasia microbiomes remained more stable. Taxa in the Family Endozoicomonadaceae, a putatively beneficial bacteria to the cnidarian holobiont (Neave et al., 2017; Rua et al., 2014; Tandon et al., 2020), were depleted in aposymbiotic, but not symbiotic, Aiptasia under antibiotic treatment. Symbiotic Aiptasia were able to restore similar microbiome communities to baseline following one week of recovery from antibiotic treatment, suggesting microbiome resilience. For both symbiotic and aposymbiotic Aiptasia, antibiotic treatment (and presumably microbiome depletion) led to downregulation of host immune pathways, but increased levels of transcription factor NF-κB, suggesting that NF-κB is involved in a non-immune aspect of microbiome depletion or antibiotic treatment (e.g., a stress response). These results imply that symbiotic and aposymbiotic cnidarians have different capacities for microbiome stability following microbiome disruption, but that they employ similar molecular mechanisms during microbiome reestablishment.

### 4.1 Antibiotic Treatment Leads to Transient Loss of Algal Symbionts

In agreement with a previous study (MacVittie et al., 2024), we observed transient changes in algal symbiont density in symbiotic Aiptasia following antibiotic treatment. In our study, R2 anemones lost colour; however, by day 7 of recovery (R7), anemones had recovered pigmentation. These results indicate that some gene expression and immune protein responses in R2 anemones could be due to changes in symbiont density. However, previous studies in Aiptasia have demonstrated that active NF-κB protein levels do not significantly increase until approximately 99% of symbionts have been lost (Mansfield et al., 2017). While different methods of symbiont density quantification were used in our study and that of Mansfield et al. (2017), R2 anemones were still visually pigmented, indicating that a high level of symbiont loss did not occur. Other metrics of host physiological response to antibiotic treatment (e.g., polyp phenotype; (Bent et al., 2021)) were not measured here, but may be important physiological responses for future studies to consider.

### 4.2 Diversity Metrics Reveal Differences in Microbiome Stability by Symbiotic State

In agreement with previous work (Bent et al., 2021), we found that antibiotic treatment significantly impacted beta diversity (here defined as dissimilarity) in both symbiotic and aposymbiotic Aiptasia. In our study, recovery from antibiotic treatment constrained diversity in community composition in symbiotic Aiptasia, suggesting that symbiotic anemones can regulate and stabilize their remaining microbiome following antibiotic treatment. However, this trend was not observed in aposymbiotic Aiptasia. Higher beta diversity measurements indicate higher variability in microbiome compositions across samples within a treatment. In cnidarians, various environmental stressors have been shown to increase microbiome beta diversity (Zaneveld et al., 2016). As suggested by the Anna Karenina principle, stochastic microbiome changes are associated with instability (Zaneveld et al., 2017), which may lead to dysbiosis (McDevitt-Irwin et al., 2017). Conversely, microbiome stability has been linked to increased resilience to heat-induced bleaching (Grottoli et al., 2018; Ziegler et al., 2017). Here, the increased microbiome stability observed in symbiotic Aiptasia during recovery may indicate higher resilience to environmental stress.

Alpha diversity metrics (inverse Simpson’s Index, Shannon Index, Richness, and Evenness) describe the diversity of the microbiome in individual samples. Here, we demonstrated that three of the four measured metrics of alpha diversity differed in symbiotic versus aposymbiotic Aiptasia across antibiotic treatments. In symbiotic Aiptasia, the inverse Simpson’s Index, the Shannon Index, and Evenness decreased before returning to baseline levels, while in aposymbiotic Aiptasia, these metrics increased before returning to baseline levels. Interestingly, previous research has demonstrated that environmental stressors (including bleaching, decreased pH, hypersalinity, and pollution) tend to increase metrics of alpha diversity (Bourne et al., 2008; Meron et al., 2011; Röthig, Ochsenkühn, et al., 2016; Ziegler et al., 2016). We demonstrate that Richness decreased following antibiotic exposure in both symbiotic and aposymbiotic Aiptasia, which corroborates previous work on microbiome depletion in anemones (MacVittie et al., 2024). Taken together, we hypothesize that aposymbiotic Aiptasia are more susceptible to microbiome shifts following disruption, which may further indicate a stressed holobiont state.

### 4.3 Microbiome Community Shifts in Aposymbiotic Aiptasia Reflect Loss of Beneficial Taxa

One striking observation from our data was the sustained depletion of Endozoicomonadaceae ASVs and overall relative abundance in aposymbiotic, but not symbiotic, Aiptasia, which lasted for at least seven days following antibiotic treatment. *Endozoicomonas* (Family Endozoicomonadaceae) has been consistently found to be an important beneficial microbial symbiont to cnidarians (Neave et al., 2017; Rua et al., 2014; Tandon et al., 2020), and depletion of this taxa is a common feature of cnidarian microbiomes following exposure to various environmental stressors (McDevitt-Irwin et al., 2017). *Endozoicomonas* has recently drawn focus for its proposed interaction with algal symbionts within cnidarian hosts. In multiple cnidarian species, including *Acropora loripes* (Gotze, Dungan, et al., 2025) and *Stylaphora pistillata* (Wada et al., 2022), host-specific strains of *Endozoicomonas* have been shown to form coral-associated microbial aggregates (CAMAs) within the host tissue. In *A. loripes*, certain *Endozoicomonas* CAMAs tightly co-localized with algal symbionts (Gotze, Dungan, et al., 2025). Additionally, different *Endozoicomonas* clades may have distinct functional roles in sugar, lipid, and phosphorous metabolism (Gotze, Tandon, et al., 2025). The association between *Endozoicomonas* and Symbiodiniaceae, and the putative clade-specific roles of *Endozoicomonas* in nutrient cycling, supports a hypothesized metabolic interaction between these two symbionts within cnidarian hosts. In our study, the depletion of Endozoicomonadaceae in aposymbiotic Aiptasia compared to the transient relative increase of these ASVs in symbiotic Aiptasia strengthens this proposed association. More specifically, our observations suggest that, because algal symbionts are housed within the symbiosome, an association between Symbiodiniaceae and Endozoicomonadaceae may buffer the bacteria from antibiotic-induced expulsion. This hypothesis could be tested using fluorescently tagged bacteria of interest and tracking their cell-level distributions over recovery (see Wada et al., 2022).

Beyond Endozoicomonadaceae, symbiotic antibiotic-treated Aiptasia were able to restore a similar bacterial community to untreated Aiptasia, indicating that symbiotic Aiptasia may employ similar strategies to microbiome regulators by maintaining a stable microbiome and relying on gene expression or physiological plasticity to withstand environmental challenges (Ziegler et al., 2019). Conversely, aposymbiotic Aiptasia failed to reestablish similar bacterial communities to untreated Aiptasia after seven days, suggesting that aposymbiotic Aiptasia use strategies analogous to microbiome conformers by altering their microbiomes in order to survive adverse conditions (Ziegler et al., 2019). While these terms (regulators and conformers) have been traditionally used for different species, it is possible that, within a species, individuals in different symbiotic states employ these same types of strategies.

In aposymbiotic anemones, we observed an increase in the relative abundance of the Families Marinobacteraceae and Moraxellaceae following antibiotic treatment. ASVs in both of these families have been associated with healthy cnidarians and holobiont tolerance to environmental stress (Cheng et al., 2023; Kvennefors et al., 2010; Ostria-Hernández et al., 2022; Randle et al., 2020; Sunagawa et al., 2009). In our aposymbiotic Aiptasia, the increase in Marinobacteraceae and Moraxellaceae and loss of Endozoicomonadaceae may thus represent a shift from one healthy microbiome to another healthy microbiome following antibiotic treatment and recovery, rather than colonization by opportunistic and non-beneficial bacteria. However, this hypothesis requires further research on the functional characteristics of the specific Marinobacteraceae and Moraxellaceae ASVs.

### 4.4 Host Regulation of Molecular Immunity May Allow for Selective Reestablishment of the Cnidarian Microbiome

Our investigation into the molecular regulation of the host transcriptome yields insights into the processes by which Aiptasia reestablishes bacterial communities following depletion. While both symbiotic states showed altered gene expression profiles following antibiotic treatment and recovery, the strength of these responses varied. Symbiotic Aiptasia exhibited greater transcriptomic shifts, signified by a higher number of differentially expressed genes (DEGs) in each treatment condition (compared to the control) versus aposymbiotic Aiptasia. This higher transcriptomic plasticity (Rivera et al., 2021) may indicate that symbiotic Aiptasia are akin to microbiome regulators in that, during environmental stress, rather than altering their microbiome, they differentially regulate gene expression pathways (Ziegler et al., 2019). Transcriptomic plasticity in combination with microbiome stability has been proposed in other cnidarians, including in the coral *Acropora cervicornis* across a depth gradient (Rodriguez-Casariego et al., 2024). Interestingly, other studies have revealed that symbiotic state often influences phenotypic responses (transcriptomic, physiological, etc.) to environmental stressors (e.g., Chei et al., 2025; Da-Anoy et al., 2025; Valadez-Ingersoll et al., 2024). For example, here and in a study in the facultatively symbiotic coral *Astrangia poculata* (Wuitchik et al., 2024), symbiotic hosts exhibited stronger transcriptomic responses to a stressor than aposymbiotic hosts (cold stress in Wuitchik et al., 2024). In contrast, under starvation stress, aposymbiotic Aiptasia had stronger transcriptomic responses than symbiotic Aiptasia (Valadez-Ingersoll et al., 2024). In a study in the facultatively symbiotic coral *Oculina arbuscula*, no differences in the magnitude of responses across symbiotic states were observed under thermal challenges (Aichelman et al., 2024). These results indicate that symbiotic state differentially modulates host responses depending on the species and the stressor. While the strength of the transcriptomic response to antibiotic treatment differed by symbiotic state, symbiotic and aposymbiotic Aiptasia showed similar patterns in specific gene expression pathways following microbiome depletion and recovery. Of note, processes relating to host metabolism were enriched following microbiome depletion, suggesting that bacterial taxa involved in nutrient cycling were depleted (Bourne et al., 2016; McDevitt-Irwin et al., 2017), and hosts must compensate for this energetic loss by increasing their metabolism. In contrast, gene expression profiles of both symbiotic and aposymbiotic Aiptasia showed downregulation of immune system pathways early in recovery from antibiotic treatment, presumably to allow for reestablishment of depleted bacteria. Our research indicated that C-Type Lectin Receptor signaling, Tumor Necrosis Factor signaling, NOD-Like Receptor signaling, the MAPK pathway, and the NF-κB pathway may be implicated in selective reestablishment of the microbiome. Of note, while immune gene expression pathways were downregulated following microbiome depletion in symbiotic and aposymbiotic Aiptasia, the levels of the active NF-κB protein increased following microbiome depletion. The Ap-NF-κB protein is regulated post-translationally (i.e., through truncation of the C-terminal inhibitory domain) (Mansfield et al., 2017). Thus, gene expression and protein activity levels of NF-κB may not necessarily correlate, and induction of the active NF-κB protein may be the result of a host stress response.

Recent work has suggested the importance of these immune system pathways for the selective uptake of compatible algal symbionts in Aiptasia (Jacobovitz et al., 2021), indicating potentially shared pathways by which the host regulates microbial and algal symbionts. While we cannot disentangle the effects of algal symbiont loss and microbiome loss on host gene expression, our work suggests future steps for understanding the molecular regulation of the cnidarian holobiont. Most studies to date exploring interactions between host immunity and the microbiome have focused on how the microbiome influences disease susceptibility (Pollock et al., 2019; Voolstra et al., 2024; Young et al., 2023). However, research in other systems, including human disease models, has revealed a great deal about the molecular interactions between the bacterial microbiome and the host immune system. For example, in microbiome establishment in mammals, the host immune system can selectively distinguish between beneficial and pathogenic bacteria (Bessman & Sonnenberg, 2016). Additionally, a wide breadth of research has investigated immunosuppression following antibiotic usage in human disease models (Ubeda & Pamer, 2012), highlighting the importance of a stable microbiome for proper immune system function. Here, we propose that, following dysbiosis of the bacterial microbiome in cnidarians, the host immune system is downregulated to allow for microbiome reestablishment, which may be an evolutionarily conserved mechanism for microbiome regulation across metazoans. Further investigation into the impact of the downregulated gene expression pathways on organismal immunity to pathogen challenge is warranted. Additionally, future research should endeavor to parse out the effect of bacterial and algal symbionts on host immunity by using axenic algal strains and microbially depleted hosts (Bove et al., 2022; Costa et al., 2021; Xiang et al., 2013).

## 5. Conclusion

This research deepens our understanding of the capacity for cnidarians to withstand disruption of their microbiomes in the context of symbiosis. In a facultatively symbiotic sea anemone, microbiome reestablishment following disruption may be regulated through similar processes in symbiotic and aposymbiotic anemones (i.e., downregulation of the immune system), but the dynamics of microbiome community composition during recovery vary by symbiotic state. For example, symbiotic anemones exhibit hallmarks of a stable microbial community that returns to baseline levels and composition after seven days, whereas aposymbiotic anemone diversity appears stochastic, and the community fails to return to the baseline state. The high stochasticity of the aposymbiotic microbiome following antibiotic treatment may be an example of the Anna Karenina principle (Zaneveld et al., 2017) in which high microbial diversity is indicative of a stressed, potentially unhealthy, holobiont. Additionally, symbiotic anemones maintain association with the putatively beneficial bacteria Endozoicomonadaceae, potentially via symbiosome evasion, which may facilitate holobiont stability throughout recovery from antibiotic treatment. This research has implications for conservation biology, as researchers rapidly respond to coral disease with antibiotic (Studivan et al., 2023) and probiotic (Ushijima et al., 2023) treatments. Additionally, it suggests a rationale for how Endozoicomonadaceae can associate with algal symbionts to increase holobiont resilience under environmental change.

## Acknowledgements

We thank the Tufts University Core Facility and the University of Texas at Austin Genomic Sequencing and Analysis Facility. We additionally thank the Boston University Marine Program, Kian Thompson, and Justin Scace for access to aquarium facilities and artificial seawater.

## Data Availability

Raw fastq files for all metabarcoding and gene expression samples are publicly available on the Sequence Read Archive BioProject PRJNA1336731. All data and code can be found here: https://github.com/mariaingersoll/Aiptasia_Microbiome.git.

