## Supplemental Tables and Figures for "Symbiotic State Affects Microbiome Recovery in a Facultatively Symbiotic Cnidarian"

### CONTENTS

**Supplementary Tables 1-6**

**Supplementary Figures 1-4**

**Table S1.** Significantly enriched Gene Ontology (GO) terms in the Biological Process (BP) category associated with immunity that were identified in the Blue module of the WGCNA containing both symbiotic and aposymbiotic samples.

| Name | Term | p.adj |
|---|---|---|
| defense response | GO:0006952; GO:0098542; GO:0051707 | 0.002835318 |
| regulation of defense response | GO:0031347 | 0.003913737 |
| response to stress | GO:0006950 | 0.004432572 |
| response to virus | GO:0009615 | 0.004432572 |
| regulation of response to stress | GO:0080134 | 0.006680102 |
| regulation of immune system process | GO:0002682 | 0.006996375 |
| immune system process | GO:0002376 | 0.00989753 |
| immune response | GO:0006955; GO:0045087; GO:0140546 | 0.01114745 |
| regulation of immune effector process | GO:0002697 | 0.01621436 |
| regulation of immune response | GO:0050776 | 0.02685909 |
| regulation of cytokine production | GO:0001817 | 0.03790314 |
| response to cytokine | GO:0034097; GO:0071345 | 0.03790314 |
| execution phase of apoptosis | GO:0097194 | 0.03907407 |
| leukocyte differentiation | GO:1903131; GO:0002521 | 0.04766559 |
| positive regulation of immune system process | GO:0002684 | 0.05216023 |
| phagocytosis | GO:0006909 | 0.06000767 |
| regulation of canonical NF-kappaB signal transduction | GO:0043122 | 0.06686703 |
| cell activation involved in immune response | GO:0002263; GO:0002366 | 0.07161406 |
| T cell receptor signaling pathway | GO:0050852 | 0.07567207 |

| | | |
|---|---|---|
| leukocyte activation | GO:0045321; GO:0046649 | 0.07161406 |
| defense response to virus | GO:0051607 | 0.07161406 |
| activation of innate immune response | GO:0002218; GO:0002221; GO:0002758 | 0.07883679 |
| immune effector process | GO:0002252 | 0.07931484 |
| T cell activation | GO:0042110 | 0.08316732 |
| positive regulation of immune response | GO:0050778 | 0.09410932 |
| mature B cell differentiation | GO:0002335 | 0.09629696 |

p.adj is the FDR *p*-value from Fisher's Exact Tests.

**Table S2.** Significantly enriched Gene Ontology (GO) terms in the Biological Process (BP) category associated with immunity identified in the Black module of the symbiotic-only WGCNA.

| Name | Term | p.adj |
|---|---|---|
| defense response | GO:0006952 | 1.126306e-05 |
| immune response | GO:0006955; GO:0045087; GO:0140546; GO:0098542 | 6.404193e-05 |
| immune system process | GO:0002376 | 8.998528e-05 |
| immune response-regulating signaling pathway | GO:0002253; GO:0002757; GO:0002764 | 0.0001963141 |
| positive regulation of immune system process | GO:0002684 | 0.0002387088 |
| positive regulation of immune response | GO:0050778 | 0.000790968 |
| regulation of immune response | GO:0050776 | 0.001462847 |
| leukocyte differentiation | GO:1903131; GO:0002521 | 0.001816461 |
| regulation of immune system process | GO:0002682 | 0.001963324 |
| antigen receptor-mediated signaling pathway | GO:0050851 | 0.002297352 |
| obsolete regulation of cell death | GO:0010941; GO:0042981; GO:0043067 | 0.004414327 |
| immune effector process | GO:0002252 | 0.006080647 |
| mature B cell differentiation | GO:0002335 | 0.006700909 |
| immune response-regulating cell surface receptor signaling pathway | GO:0002768 | 0.006700909 |
| T cell receptor signaling pathway | GO:0050852 | 0.007644907 |
| negative regulation of programmed cell death | GO:0043066; GO:0043069; GO:0060548 | 0.0100702 |
| regulation of cytokine production | GO:0001817 | 0.01137308 |
| response to virus | GO:0009615 | 0.01171845 |
| response to cytokine | GO:0034097; GO:0071345 | 0.01336926 |
| B cell differentiation | GO:0030183 | 0.0133862 |
| regulation of leukocyte differentiation | GO:1902105; GO:1902107; GO:1903708 | 0.016138 |
| leukocyte activation | GO:0045321; GO:0046649 | 0.016805 |
| lymphocyte differentiation | GO:0030098 | 0.01714824 |
| cytokine-mediated signaling pathway | GO:0019221 | 0.01762023 |

| | | |
|---|---|---|
| immune response-<br>activating cell surface receptor signaling pathway | GO:0002429 | 0.02339626 |
| positive regulation of NF-kappaB transcription factor activity | GO:0051092 | 0.05664149 |
| regulation of interleukin-2 production | GO:0032663 | 0.0569638 |
| regulation of B cell activation | GO:0050864 | 0.0569638 |
| regulation of canonical NF-kappaB signal transduction | GO:0043122 | 0.06086602 |

p.adj is the FDR *p*-value from Fisher's Exact Tests.

**Table S3.** Significantly enriched Gene Ontology (GO) terms in the Biological Process (BP) category associated with immunity identified in the Blue module of the aposymbiotic-only WGCNA.

| **Name** | **Term** | **p.adj** |
|---|---|---|
| immune system process | GO:0002376 | 0.001471418 |
| regulation of myeloid leukocyte differentiation | GO:0002761; GO:0002763; GO:0045639 | 0.02349169 |
| defense response | GO:0006952 | 0.02853583 |
| immune response | GO:0006955; GO:0045087; GO:0140546; GO:0098542 | 0.02853583 |
| regulation of immune system process | GO:0002682 | 0.05254129 |
| obsolete regulation of necrotic cell death | GO:0010939 | 0.05787996 |
| response to tumor necrosis factor | GO:0034612 | 0.05787996 |
| execution phase of apoptosis | GO:0097194 | 0.07453446 |

p.adj is the FDR *p*-value from Fisher's Exact Tests.

**Table S4.** Raw and filtered read counts for samples used in 16S profiling. Sample Alias refers to the library ID found in the SRA data.

| Sample | Sample Alias | Treatment | Symbiotic State | Raw | Filtered |
|---|---|---|---|---|---|
| CS7 | CS7 | C | Sym | 74740 | 60483 |
| CS8 | CS8 | C | Sym | 75079 | 60213 |
| CS9 | CS9 | C | Sym | 61322 | 48494 |
| CS10 | CS10 | C | Sym | 32428 | 24830 |
| CS11 | CS11 | C | Sym | 75250 | 56300 |
| CS12 | CS12 | C | Sym | 64231 | 48172 |
| R0S8 | AS8 | R0 | Sym | 39256 | 25599 |
| R0S9 | AS9 | R0 | Sym | 37385 | 21241 |
| R0S10 | AS10 | R0 | Sym | 35981 | 20550 |
| R0S11 | AS11 | R0 | Sym | 18497 | 8572 |
| R0S12 | AS12 | R0 | Sym | 21768 | 13810 |
| R0S13 | AS13 | R0 | Sym | 68791 | 48756 |
| R2S7 | PS7 | R2 | Sym | 25193 | 13867 |
| R2S8 | PS8 | R2 | Sym | 21959 | 10149 |
| R2S9 | PS9 | R2 | Sym | 63923 | 53537 |
| R2S10 | PS10 | R2 | Sym | 18603 | 8680 |
| R2S11 | PS11 | R2 | Sym | 60075 | 40664 |
| R2S12 | PS12 | R2 | Sym | 34264 | 20962 |
| R7S8 | RS8 | R7 | Sym | 36019 | 21577 |
| R7S9 | RS9 | R7 | Sym | 67581 | 48831 |
| R7S10 | RS10 | R7 | Sym | 59545 | 42374 |
| R7S11 | RS11 | R7 | Sym | 49900 | 33647 |
| R7S12 | RS12 | R7 | Sym | 37258 | 23706 |
| R7S13 | RS13 | R7 | Sym | 36123 | 24152 |
| R7S14 | RS14 | R7 | Sym | 50939 | 31606 |
| CA7 | CA7 | C | Apo | 77573 | 63283 |
| CA8 | CA8 | C | Apo | 53451 | 42320 |
| CA9 | CA9 | C | Apo | 57081 | 37395 |
| CA10 | CA10 | C | Apo | 64471 | 53172 |
| CA11 | CA11 | C | Apo | 53058 | 40612 |
| CA12 | CA12 | C | Apo | 108221 | 84324 |
| R0A2 | AA2 | R0 | Apo | 26838 | 15166 |
| R0A3 | AA3 | R0 | Apo | 19351 | 10348 |
| R0A9 | AA9 | R0 | Apo | 14633 | 7625 |
| R0A10 | AA10 | R0 | Apo | 21983 | 7509 |
| R0A11 | AA11 | R0 | Apo | 13078 | 8557 |
| R0A12 | AA12 | R0 | Apo | 22212 | 12914 |
| R2A7 | PA7 | R2 | Apo | 58808 | 43475 |
| R2A8 | PA8 | R2 | Apo | 40700 | 27074 |

| | | | | | |
|---|---|---|---|---|---|
| R2A10 | PA10 | R2 | Apo | 56330 | 40143 |
| R2A11 | PA11 | R2 | Apo | 45231 | 33800 |
| R2A12 | PA12 | R2 | Apo | 36328 | 23000 |
| R7A7 | RA7 | R7 | Apo | 87216 | 67753 |
| R7A8 | RA8 | R7 | Apo | 65256 | 52827 |
| R7A9 | RA9 | R7 | Apo | 48640 | 38941 |
| R7A10 | RA10 | R7 | Apo | 55922 | 45696 |
| R7A11 | RA11 | R7 | Apo | 12017 | 9775 |
| R7A12 | RA12 | R7 | Apo | 46261 | 33920 |

Sym = symbiotic; Apo = aposymbiotic.

**Table S5.** Number of differentially expressed genes (DEGs; FDR *p*-value of <0.05) in Aiptasia for all pairwise comparisons between treatments within symbiotic states or between symbiotic states within treatments.

| Comparison | DEGs |
|---|---|
| R0A:CA | 814 |
| R2A:CA | 958 |
| R7A:CA | 342 |
| R0A:R7A | 924 |
| R2A:R7A | 632 |
| R0A:R2A | 805 |
| R0S:CS | 1000 |
| R2S:CS | 983 |
| R7S:CS | 1347 |
| R0S:R7S | 1626 |
| R2S:R7S | 1014 |
| R0S:R2S | 946 |
| CS:CA | 2374 |
| R0S:R0A | 2520 |
| R2S:R2A | 1968 |
| R7S:R7A | 1834 |

CS = control symbiotic; R0S = R0 symbiotic; R2S = R2 symbiotic; R7S = R7 symbiotic; CA = control aposymbiotic; R0A = R0 aposymbiotic; R2A = R2 aposymbiotic; R7A = R7 aposymbiotic.

**Table S6.** Delta rank values of enriched Gene Ontology (GO) terms in the Biological Process (BP) category were compared across each treatment condition relative to the control (within symbiotic state). Terms with positive values are significantly enriched in the treatment, and terms with negative values are significantly enriched in the control. NA indicates that a term was not significantly enriched in that comparison.

| GO Term | SR7C | AR7C | SR2C | AR2C | AR0C | SR0C |
|---|---|---|---|---|---|---|
| actin filament capping | 2476 | -1922 | NA | NA | -1905 | NA |
| actin filament-based process | 922 | NA | NA | NA | -768 | 982 |
| alpha-amino acid biosynthetic process | NA | NA | NA | 1046 | 1223 | NA |
| alpha-amino acid metabolic process | NA | NA | 656 | 788 | 1093 | NA |
| amine metabolic process | 1214 | NA | NA | NA | 1165 | NA |
| aminoglycan catabolic process | NA | NA | NA | 1460 | NA | -1449 |
| aminoglycan metabolic process | NA | 1178 | NA | 1010 | NA | NA |
| anatomical structure morphogenesis | 386 | -332 | NA | -524 | -298 | 418 |
| anion transport | 987 | NA | 738 | NA | NA | NA |
| axonemal dynein complex assembly | NA | -2093 | NA | -2092 | NA | NA |
| calcium ion transmembrane transport | NA | NA | NA | -834 | -725 | -780 |
| carbohydrate metabolic process | NA | NA | NA | 495 | 368 | NA |
| cardiac muscle cell action potential | 2887 | NA | NA | NA | -2778 | NA |
| cardiac septum development | NA | -1685 | NA | -1775 | -2066 | NA |
| cation transport | 783 | NA | NA | NA | NA | -489 |
| cell communication | 672 | NA | NA | NA | NA | -529 |
| cell cycle | NA | -672 | 382 | NA | -687 | NA |
| cell cycle process | NA | -626 | NA | NA | -652 | NA |
| cell division | NA | -718 | NA | NA | -802 | NA |
| cell fate commitment involved in formation of primary germ layer | NA | NA | NA | -2768 | -2469 | NA |
| cell killing | NA | 1948 | 1599 | 1965 | NA | NA |
| cell motility | NA | NA | NA | -521 | -752 | NA |
| cell part morphogenesis | NA | -937 | NA | -1009 | -823 | NA |
| cell projection assembly | NA | -1031 | NA | NA | NA | 1455 |

| | | | | | | |
|---|---|---|---|---|---|---|
| cell projection organization | NA | -891 | NA | -870 | -817 | 511 |
| cell redox homeostasis | NA | 1272 | NA | NA | NA | -2134 |
| cell surface receptor signaling pathway | 338 | NA | -277 | -566 | -371 | NA |
| cellular aldehyde metabolic process | NA | NA | NA | 2285 | 1457 | NA |
| cellular amino acid biosynthetic process | NA | NA | NA | 990 | 1119 | NA |
| cellular amino acid metabolic process | NA | NA | NA | 521 | 709 | NA |
| cellular component assembly | NA | NA | 318 | NA | -456 | NA |
| cellular component assembly involved in morphogenesis | NA | NA | NA | -806 | -675 | 1157 |
| cellular component biogenesis | NA | NA | NA | 802 | 1061 | NA |
| cellular component morphogenesis | NA | -741 | NA | -734 | -686 | NA |
| cellular component movement | NA | -590 | NA | -654 | -862 | NA |
| cellular hormone metabolic process | NA | NA | NA | 1137 | 1614 | NA |
| cellular lipid metabolic process | NA | NA | 585 | 506 | 394 | NA |
| cellular modified amino acid biosynthetic process | NA | NA | NA | 1317 | 1122 | NA |
| cellular modified amino acid metabolic process | NA | NA | 819 | 943 | 1250 | NA |
| cellular response to DNA damage stimulus | -521 | NA | NA | NA | NA | 445 |
| cGMP metabolic process | 2158 | NA | NA | NA | -1965 | NA |
| chloride transport | 2480 | NA | 1492 | NA | NA | NA |
| chromatin organization | NA | -531 | NA | NA | -460 | 479 |
| chromosome organization | -474 | -494 | NA | NA | -343 | NA |
| chromosome segregation | NA | NA | 1146 | 1503 | NA | NA |
| cilium morphogenesis | NA | -1597 | NA | -1364 | -1213 | NA |
| cilium movement | NA | -2086 | NA | -2037 | -1363 | NA |
| cilium organization | NA | -1109 | NA | -1091 | NA | 1124 |
| cofactor biosynthetic process | NA | NA | NA | 929 | 1205 | NA |
| cofactor metabolic process | NA | NA | 827 | 877 | 1219 | 1012 |

| | | | | | | |
|---|---|---|---|---|---|---|
| convergent extension | NA | -1790 | NA | -1624 | -1379 | NA |
| covalent chromatin modification | NA | -622 | NA | -624 | NA | 804 |
| Cytokinesis | NA | -1051 | NA | NA | -973 | 1071 |
| cytoskeleton organization | NA | -732 | NA | -623 | -921 | 893 |
| Death | NA | NA | -426 | -402 | NA | NA |
| defense response | NA | NA | -785 | -539 | NA | NA |
| defense response to other organism | NA | NA | -1097 | -1059 | NA | NA |
| Digestion | NA | 1933 | NA | NA | -1287 | -1804 |
| DNA integration | -740 | NA | -609 | NA | NA | -1105 |
| DNA metabolic process | -767 | NA | NA | NA | NA | -458 |
| DNA replication | -979 | NA | NA | NA | NA | -1102 |
| electron transport chain | NA | NA | NA | 1786 | 1409 | NA |
| embryonic heart tube morphogenesis | NA | -1277 | NA | -1655 | NA | NA |
| endosomal transport | NA | -788 | -907 | -1063 | NA | NA |
| epithelial cilium movement | NA | -1950 | NA | -1493 | NA | NA |
| establishment of localization in cell | -405 | -342 | -338 | -518 | NA | NA |
| establishment of mitotic spindle localization | NA | -1760 | NA | NA | -2441 | NA |
| establishment of planar polarity of embryonic epithelium | NA | NA | -2630 | NA | -2342 | -2500 |
| establishment of protein localization | -650 | NA | -321 | NA | NA | 354 |
| establishment of spindle localization | NA | -1864 | NA | -1816 | -2166 | NA |
| establishment or maintenance of cell polarity | NA | -895 | NA | -1338 | -1545 | NA |
| extracellular matrix disassembly | NA | 1314 | NA | NA | NA | -1447 |
| extracellular structure organization | NA | NA | NA | NA | -629 | -1049 |
| fatty acid beta-oxidation | NA | NA | 1939 | 1979 | NA | NA |
| fatty acid catabolic process | NA | NA | 1844 | 1322 | NA | NA |
| fatty acid metabolic process | NA | NA | 1196 | 854 | 722 | NA |

| | | | | | | |
|---|---|---|---|---|---|---|
| G-protein coupled receptor signaling pathway | 626 | NA | NA | NA | NA | -652 |
| glutathione metabolic process | NA | NA | NA | 1468 | 1325 | NA |
| glycosyl compound biosynthetic process | NA | NA | NA | 970 | 1300 | NA |
| heme metabolic process | NA | NA | NA | 1896 | 1966 | NA |
| histone H3-K4 trimethylation | NA | NA | NA | -3347 | -2584 | NA |
| hydrogen transport | NA | NA | NA | 1033 | 1229 | NA |
| immune effector process | NA | NA | -885 | -1015 | NA | NA |
| immune response | NA | NA | -993 | -640 | NA | NA |
| immune system process | NA | NA | -799 | -647 | -514 | NA |
| inner ear receptor stereocilium organization | NA | -1549 | NA | -1665 | -1766 | NA |
| inorganic anion transport | 2043 | NA | 1782 | NA | NA | NA |
| intracellular protein transport | -763 | NA | NA | NA | NA | 576 |
| intracellular signal transduction | NA | -401 | NA | -474 | -441 | 398 |
| ion transmembrane transport | 872 | NA | NA | NA | NA | -646 |
| isoprenoid metabolic process | NA | NA | 1277 | NA | 1297 | NA |
| kidney morphogenesis | NA | -1654 | NA | -1927 | NA | NA |
| leukocyte activation | NA | NA | -994 | -1092 | NA | NA |
| leukocyte differentiation | NA | NA | -1255 | -1060 | NA | NA |
| leukotriene metabolic process | NA | NA | 2814 | 2420 | NA | NA |
| lipid biosynthetic process | NA | NA | 555 | 661 | 497 | NA |
| lipid metabolic process | NA | NA | 553 | 375 | 332 | NA |
| Locomotion | NA | NA | NA | -546 | -682 | NA |
| long-chain fatty acid metabolic process | NA | NA | 1730 | 1321 | NA | NA |
| lung cell differentiation | NA | -2491 | NA | -2638 | NA | NA |
| lung development | NA | -1091 | -1217 | -1319 | NA | NA |
| lymphocyte differentiation | NA | NA | -1449 | -1074 | NA | NA |
| maintenance of organ identity | NA | -1765 | -2036 | -1938 | -1760 | NA |

| | | | | | | |
|---|---|---|---|---|---|---|
| meiotic nuclear division | NA | NA | 1615 | 1008 | NA | NA |
| metal ion transport | 967 | NA | NA | NA | NA | -504 |
| microtubule cytoskeleton organization | NA | -873 | NA | NA | -841 | 852 |
| microtubule-based movement | NA | -1267 | NA | -1214 | -1202 | NA |
| microtubule-based process | NA | -952 | NA | -627 | -1023 | 491 |
| mitotic cell cycle | NA | NA | 1219 | 835 | NA | NA |
| mitotic spindle elongation | NA | NA | NA | 3234 | 2602 | NA |
| modification of morphology or physiology of other organism | NA | 1374 | NA | 1229 | NA | -1267 |
| monocarboxylic acid catabolic process | NA | NA | 1876 | 1351 | NA | NA |
| monocarboxylic acid metabolic process | NA | NA | 1028 | 697 | 624 | NA |
| monosaccharide metabolic process | NA | NA | 1334 | 825 | 985 | 916 |
| mRNA metabolic process | -609 | NA | NA | NA | NA | 885 |
| multicellular organismal development | 514 | NA | NA | NA | NA | 571 |
| myeloid leukocyte activation | NA | NA | NA | -1730 | -1674 | NA |
| NADP metabolic process | NA | NA | 2266 | 2173 | NA | NA |
| negative regulation of biosynthetic process | NA | NA | NA | -347 | NA | 707 |
| negative regulation of cell proliferation | NA | -589 | NA | -860 | NA | NA |
| negative regulation of coagulation | NA | 2829 | NA | 1657 | NA | -2002 |
| negative regulation of developmental process | NA | NA | NA | -643 | -874 | NA |
| negative regulation of inclusion body assembly | NA | NA | -2945 | -2642 | NA | NA |
| negative regulation of metabolic process | NA | -327 | NA | -467 | NA | 718 |
| negative regulation of protein metabolic process | NA | NA | NA | -728 | NA | 624 |
| negative regulation of response to stimulus | NA | -364 | NA | -558 | NA | NA |

| | | | | | | |
|---|---|---|---|---|---|---|
| neuropeptide signaling pathway | 990 | NA | NA | NA | -601 | NA |
| nucleic acid phosphodiester bond hydrolysis | -692 | NA | -562 | NA | NA | -688 |
| nucleoside monophosphate biosynthetic process | -1390 | NA | NA | 1190 | 1494 | NA |
| one-carbon metabolic process | NA | 1747 | NA | NA | 2266 | NA |
| organ development | NA | NA | NA | -395 | -343 | NA |
| organelle assembly | NA | -615 | NA | NA | -543 | 938 |
| organelle fission | NA | -666 | 586 | NA | -663 | NA |
| organelle localization | NA | NA | NA | -1368 | -1364 | NA |
| organic acid metabolic process | NA | NA | 647 | 669 | 701 | NA |
| organic cation transport | 2504 | NA | 2277 | NA | NA | NA |
| organic cyclic compound catabolic process | NA | -356 | NA | NA | NA | 396 |
| organonitrogen compound biosynthetic process | NA | 465 | 513 | 708 | 785 | NA |
| organonitrogen compound metabolic process | NA | NA | 350 | 254 | 268 | NA |
| Ossification | NA | NA | -1284 | NA | NA | -1357 |
| oxidation-reduction process | 479 | 361 | 1011 | 844 | 1041 | 677 |
| oxidoreduction coenzyme metabolic process | NA | NA | 2029 | NA | NA | 1605 |
| Pathogenesis | NA | NA | 2166 | 2059 | NA | NA |
| pentose-phosphate shunt | NA | NA | 2751 | 2629 | NA | NA |
| peptide catabolic process | NA | NA | 2137 | 2087 | NA | NA |
| peptide metabolic process | NA | NA | NA | 1120 | 1140 | NA |
| peptidyl-amino acid modification | NA | NA | NA | -488 | -354 | 489 |
| peptidyl-lysine modification | NA | NA | NA | -885 | NA | 1032 |
| peptidyl-proline hydroxylation | 2840 | NA | 2625 | NA | NA | NA |
| peptidyl-proline modification | NA | NA | 1867 | NA | NA | 1388 |

| | | | | | | |
|---|---|---|---|---|---|---|
| peptidyl-tyrosine dephosphorylation | NA | NA | -1362 | -1222 | -1111 | NA |
| peptidyl-tyrosine phosphorylation | NA | NA | NA | -1012 | -716 | NA |
| Phosphorylation | NA | -440 | NA | -481 | -358 | NA |
| photoreceptor cell maintenance | NA | -1330 | -1326 | -1541 | -1242 | NA |
| positive regulation of blood coagulation | NA | NA | -2834 | NA | NA | -2293 |
| positive regulation of defense response | NA | NA | -1355 | -1088 | NA | NA |
| positive regulation of immune response | NA | NA | -1491 | -1084 | NA | NA |
| positive regulation of immune system process | NA | NA | -913 | -702 | NA | NA |
| positive regulation of metabolic process | NA | NA | -301 | NA | -245 | NA |
| positive regulation of molecular function | NA | NA | NA | -387 | -307 | NA |
| positive regulation of response to stimulus | NA | NA | -465 | -568 | -354 | NA |
| positive regulation of transport | NA | NA | NA | -553 | -583 | NA |
| potassium ion transport | 1493 | NA | NA | NA | NA | -1114 |
| primary alcohol metabolic process | NA | NA | NA | 1695 | 2822 | 1773 |
| protein autoubiquitination | 1414 | NA | -1220 | -1617 | NA | NA |
| protein dephosphorylation | NA | NA | -907 | -1130 | NA | NA |
| protein folding | -1488 | 1124 | NA | 1082 | NA | NA |
| protein hydroxylation | 2250 | NA | 1931 | NA | NA | 2084 |
| protein localization to paranode region of axon | NA | -3166 | NA | NA | -2905 | NA |
| protein modification by small protein conjugation or removal | NA | -625 | -633 | -811 | -355 | NA |
| protein modification by small protein removal | NA | NA | NA | -1035 | -1004 | NA |
| protein-DNA complex assembly | NA | NA | 1445 | 1226 | NA | NA |
| Proteolysis | -564 | 757 | -494 | NA | NA | -658 |

| | | | | | | |
|---|---|---|---|---|---|---|
| pteridine-containing compound metabolic process | NA | 1563 | NA | NA | 2155 | NA |
| regulation of actin filament depolymerization | 2434 | NA | NA | NA | -1559 | NA |
| regulation of anatomical structure morphogenesis | NA | NA | NA | -529 | -672 | NA |
| regulation of bone remodeling | NA | NA | -2426 | -2502 | NA | NA |
| regulation of canonical Wnt signaling pathway | NA | -1049 | NA | -916 | -949 | NA |
| regulation of catabolic process | NA | -408 | NA | -528 | -358 | 483 |
| regulation of cell communication | NA | -324 | -289 | -543 | -412 | NA |
| regulation of cell cycle | NA | -569 | NA | -446 | -444 | 425 |
| regulation of cell differentiation | NA | NA | NA | -607 | -504 | NA |
| regulation of cell division | NA | NA | 1170 | NA | NA | 1112 |
| regulation of cell proliferation | 424 | NA | NA | -502 | -375 | NA |
| regulation of cellular component organization | NA | -278 | NA | -422 | -469 | NA |
| regulation of cellular localization | NA | NA | NA | -606 | -424 | NA |
| regulation of chromatin organization | NA | NA | NA | -1367 | NA | 1670 |
| regulation of chromosome organization | NA | -980 | NA | NA | NA | 1176 |
| regulation of cyclic nucleotide metabolic process | NA | NA | -906 | -1236 | -1080 | -1250 |
| regulation of developmental process | NA | -286 | NA | -553 | -553 | NA |
| regulation of epithelial to mesenchymal transition | NA | 1816 | NA | 1495 | NA | NA |
| regulation of immune effector process | NA | NA | -1403 | -1147 | NA | NA |
| regulation of immune response | NA | NA | -973 | -855 | NA | NA |
| regulation of immune system process | NA | NA | -518 | -726 | -437 | NA |
| regulation of innate immune response | NA | NA | -1260 | -921 | NA | NA |
| regulation of interleukin-2 production | NA | NA | -2429 | -2331 | NA | NA |

| | | | | | | |
|---|---|---|---|---|---|---|
| regulation of intracellular protein transport | NA | -881 | NA | -838 | NA | 1109 |
| regulation of intracellular signal transduction | NA | NA | NA | -541 | -413 | NA |
| regulation of intracellular transport | NA | -759 | NA | -832 | NA | 1206 |
| regulation of ion transport | 1151 | NA | NA | NA | NA | -654 |
| regulation of localization | 511 | NA | NA | -408 | -373 | NA |
| regulation of metal ion transport | 1161 | NA | NA | NA | -938 | NA |
| regulation of mitotic cell cycle | NA | -807 | NA | -680 | -703 | NA |
| regulation of molecular function | NA | -307 | NA | -368 | -311 | NA |
| regulation of multicellular organismal process | 343 | NA | NA | -422 | -356 | NA |
| regulation of myeloid cell differentiation | NA | NA | NA | -1392 | -1139 | NA |
| regulation of neural precursor cell proliferation | 1329 | NA | NA | -1229 | NA | NA |
| regulation of nucleocytoplasmic transport | NA | -914 | NA | -933 | NA | 1332 |
| regulation of nucleoside metabolic process | NA | -453 | NA | -701 | -519 | 565 |
| regulation of nucleotide metabolic process | NA | -554 | NA | -845 | -671 | NA |
| regulation of organ morphogenesis | NA | NA | NA | -940 | -888 | NA |
| regulation of organelle organization | NA | -513 | NA | NA | NA | 672 |
| regulation of osteoblast proliferation | NA | -2159 | -2595 | -1893 | NA | NA |
| regulation of phosphorus metabolic process | NA | -336 | NA | -572 | -477 | NA |
| regulation of planar cell polarity pathway involved in neural tube closure | NA | -2371 | NA | -2231 | -2068 | NA |

| | | | | | | |
|---|---|---|---|---|---|---|
| regulation of protein complex disassembly | 1460 | NA | NA | NA | -1289 | NA |
| regulation of protein depolymerization | 1780 | NA | NA | NA | -1298 | NA |
| regulation of protein metabolic process | NA | NA | -413 | -323 | NA | NA |
| regulation of Ras GTPase activity | NA | -657 | NA | -789 | -765 | NA |
| regulation of response to stress | NA | NA | -474 | -459 | NA | NA |
| regulation of Rho protein signal transduction | NA | NA | NA | -1205 | -1095 | NA |
| regulation of small GTPase mediated signal transduction | NA | NA | NA | -978 | -895 | NA |
| regulation of stem cell differentiation | 1499 | 1199 | NA | NA | NA | NA |
| regulation of transport | 470 | NA | NA | -503 | -332 | NA |
| regulation of vasodilation | NA | NA | NA | NA | -1632 | -2042 |
| response to biotic stimulus | NA | NA | -751 | -663 | -492 | -604 |
| response to external stimulus | NA | NA | NA | -471 | -342 | NA |
| response to mechanical stimulus | 1224 | NA | NA | -708 | -852 | NA |
| response to other organism | NA | NA | -962 | -834 | -566 | -660 |
| response to virus | NA | NA | -982 | -986 | NA | NA |
| retinol metabolic process | NA | NA | NA | NA | 2882 | 2056 |
| ribonucleoprotein complex biogenesis | NA | NA | NA | 909 | 1054 | NA |
| RNA biosynthetic process | NA | -326 | NA | -257 | NA | 537 |
| RNA processing | -528 | NA | NA | NA | NA | 778 |
| RNA-dependent DNA replication | -875 | NA | -827 | NA | NA | -1471 |
| rRNA metabolic process | NA | NA | NA | 895 | 909 | NA |
| single organism signaling | 889 | NA | NA | NA | NA | -658 |
| single-organism biosynthetic process | NA | NA | 513 | 683 | 666 | 368 |
| single-organism organelle organization | NA | -726 | NA | -407 | -620 | 561 |
| small molecule biosynthetic process | NA | NA | 579 | 847 | 832 | NA |
| small molecule catabolic process | NA | NA | 853 | 639 | 894 | 800 |
| sodium ion transport | 1242 | 855 | NA | 1082 | NA | NA |

| | | | | | | |
|---|---|---|---|---|---|---|
| specification of symmetry | NA | -1252 | NA | -1330 | -919 | NA |
| steroid biosynthetic process | NA | NA | 1246 | NA | 881 | 1568 |
| steroid metabolic process | NA | NA | 851 | NA | NA | 814 |
| sulfur amino acid metabolic process | NA | NA | 1246 | NA | 1312 | NA |
| sulfur compound metabolic process | NA | NA | 725 | 720 | 778 | NA |
| system process | 618 | NA | NA | NA | NA | -414 |
| tissue morphogenesis | NA | NA | NA | -887 | -559 | NA |
| Translation | -872 | 921 | -1389 | 1022 | 1135 | -1167 |
| translational elongation | NA | 1624 | -1889 | NA | 2333 | NA |
| translational initiation | -1852 | 967 | NA | 1146 | NA | NA |
| tRNA aminoacylation for protein translation | -3855 | -1735 | -3262 | -2255 | -2259 | -2737 |
| tRNA metabolic process | -1481 | -984 | NA | -828 | NA | NA |
| trypsinogen activation | 3736 | NA | 3264 | NA | NA | 2827 |
| unsaturated fatty acid metabolic process | NA | NA | 1708 | 1466 | NA | NA |
| vesicle-mediated transport | NA | NA | -351 | -344 | -343 | NA |
| vitamin metabolic process | NA | NA | NA | NA | 1421 | 1262 |
| Wnt signaling pathway | NA | -593 | NA | -633 | -714 | NA |

SR7C=symbiotic R7 to symbiotic control; AR7C=aposymbiotic R7 to aposymbiotic control; SR2C=symbiotic R2 to symbiotic control; AR2C=aposymbiotic R2 to aposymbiotic control; AR0C=aposymbiotic R0 to aposymbiotic control; SR0C=symbiotic R0 to symbiotic control.

**FIGURE S1**

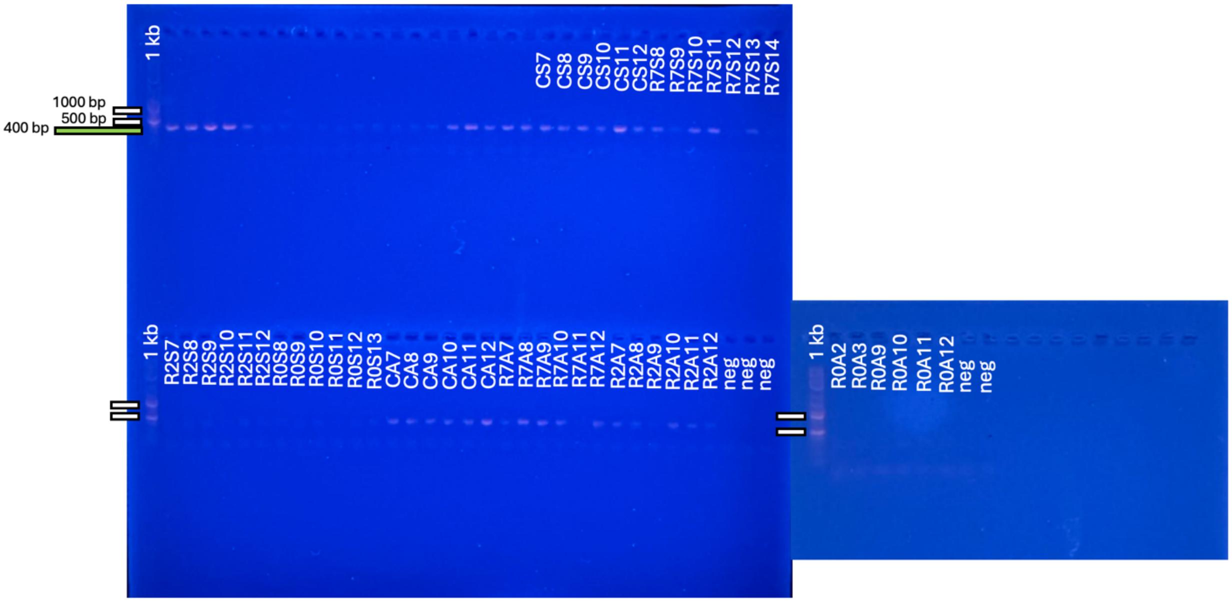


**Figure S1. Amplification of 16S gene from symbiotic and aposymbiotic Aiptasia across antibiotic treatments.** The 16S rRNA gene was amplified from each sample and PCR products from each sample were purified and visualized on two 1% agarose gels. Lanes without labels were samples not relevant to the current experiment, but they serve as a size marker for the correct PCR product. Included in the gels are 1 kb ladders (1 kb). Negative controls (neg) are included in each gel and contain molecular grade water instead of DNA as template and were sequenced to bioinformatically remove contaminants. CS = control symbiotic; R7S = recovery7 symbiotic; R2S = recovery2 symbiotic; R0S = recovery0 symbiotic; CA = control aposymbiotic; R7A = recovery7 aposymbiotic; R2A = recovery2 aposymbiotic; R0A = recovery0 aposymbiotic. White bars to the left of the gels indicate the 1000 bp and 500 bp markers on the ladder. Green bars to the left of the gel indicate ~400 bp, the approximate size of the PCR product.

**FIGURE S2**

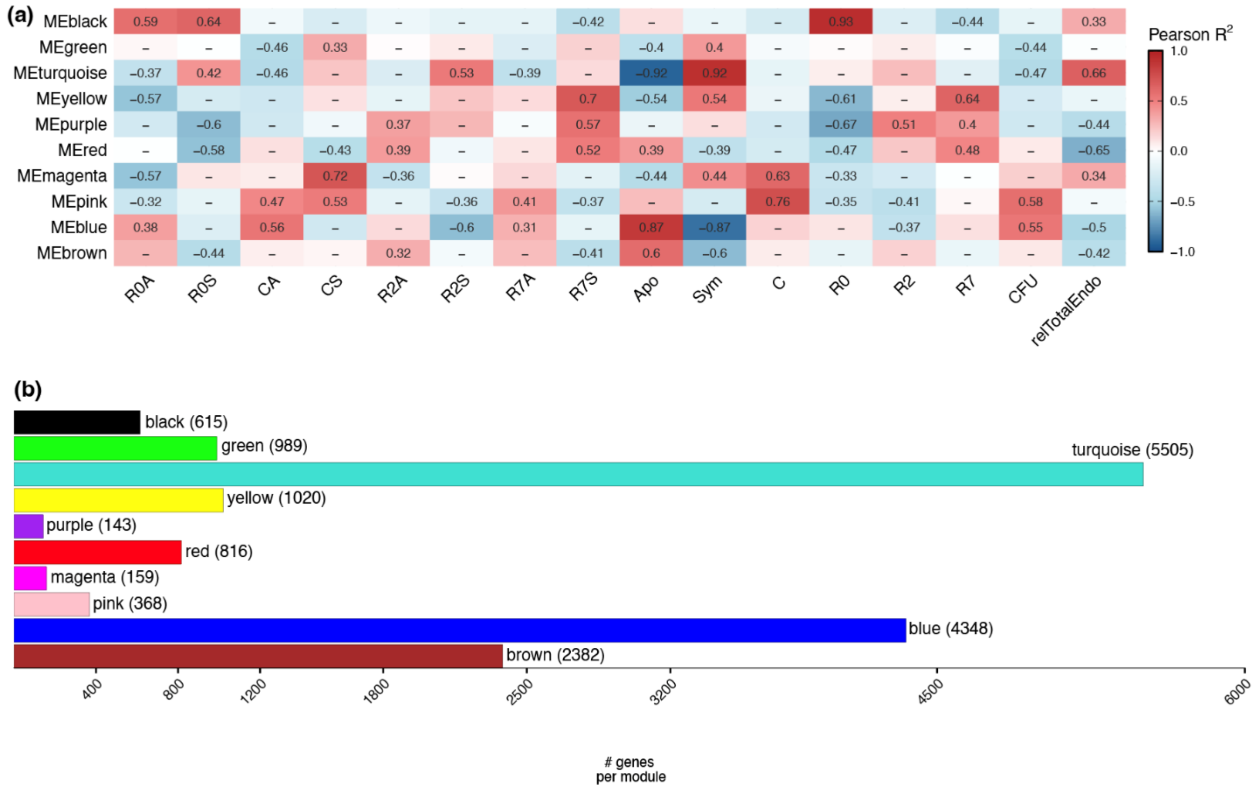


**Figure S2. Modules identified for WGCNA from the combined model with symbiotic and aposymbiotic Aiptasia. (a)** Experimental treatments and microbiome traits of Aiptasia were correlated with module eigengene expression of modules identified through WGCNA. Colours within the heat map correspond to directionality of the Pearson $R^2$ value, in which positive correlations are red and negative correlations are blue. Cells that contain values identify significant relationships between a treatment/trait and a module's eigengene expression. CFU = normalized total Colony Forming Units (**Fig. 1a**); relTotalEndo = relative abundance of the summed counts for Endozoicomonadaceae ASVs (**Fig. 3b**). **(b)** The number of genes belonging to each identified module are identified in a bar graph in which colours correspond to the names of the modules. R0A = R0 aposymbiotic; R0S = R0 symbiotic; CA = control aposymbiotic; CS = control symbiotic; R2A = R2 aposymbiotic; R2S = R2 symbiotic; R7A = R7 aposymbiotic; R7S = R7 symbiotic.

### FIGURE S3

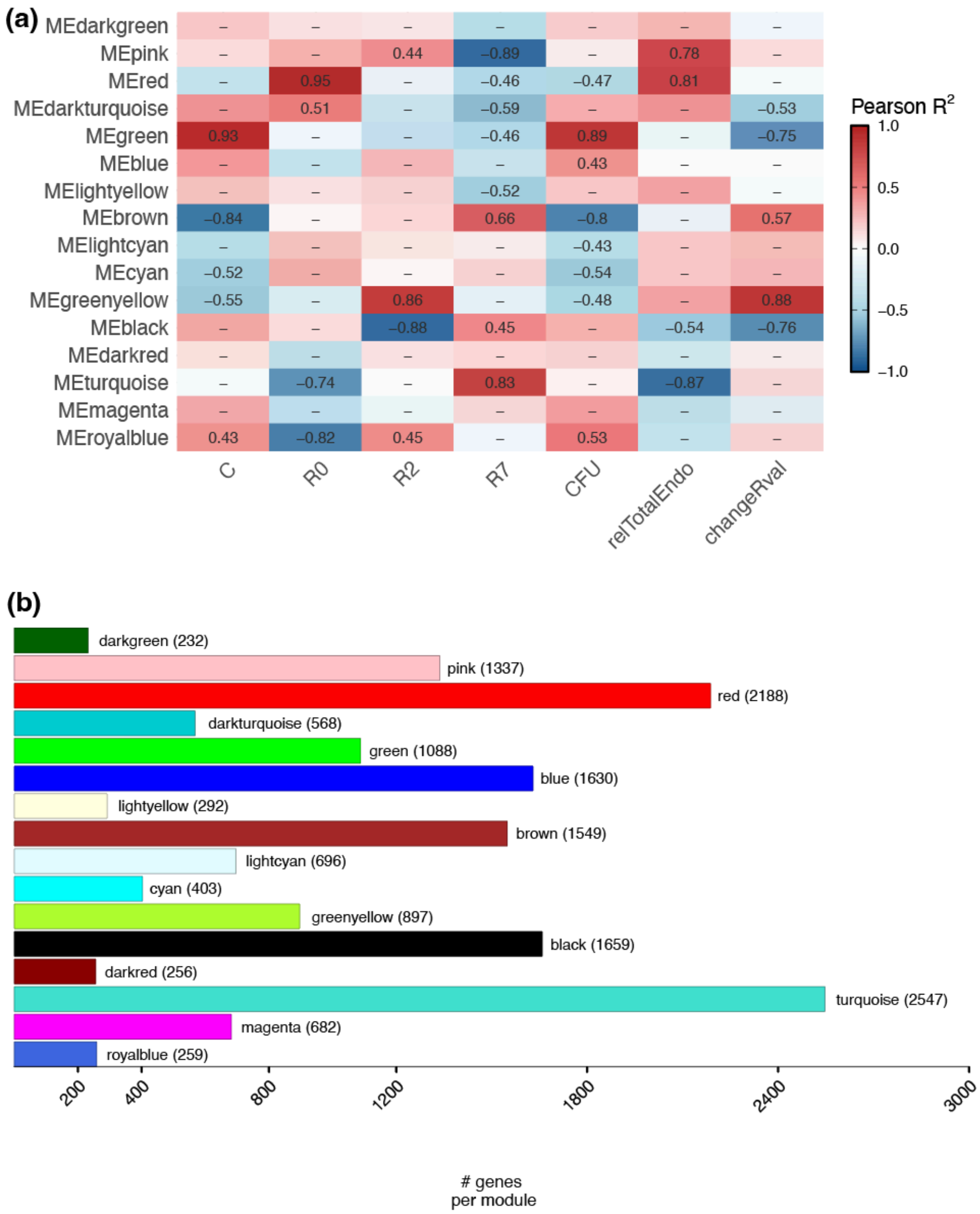


**Figure S3. WGCNA modules identified for symbiotic Aiptasia. (a)** Experimental treatments and microbiome traits of symbiotic Aiptasia were correlated with module eigengene expression of modules identified through WGCNA. Colours within the heat map correspond to directionality of the Pearson $R^2$ value, in which positive correlations are red and negative correlations are blue. Cells that contain values identify significant relationships between a treatment/trait and a module's eigengene expression. CFU = normalized total Colony Forming Units (**Fig. 1a**); relTotalEndo = relative abundance of the summed counts for Endozoicomonadaceae ASVs (**Fig. 3b**); changeRval = change in red channel intensity (**Fig. 1b**). **(b)** The number of genes belonging to each identified module are identified in a bar graph in which colours correspond to the names of the modules.

**FIGURE S4**

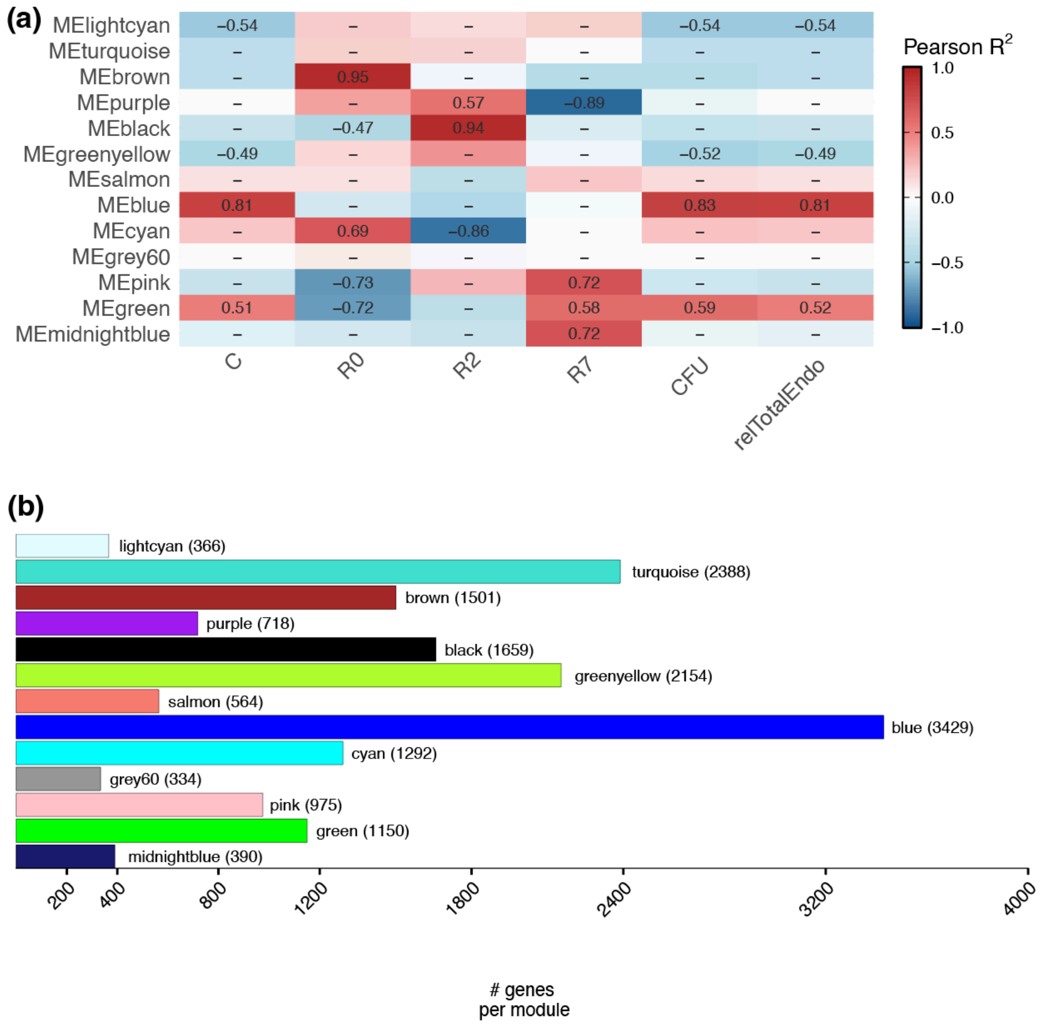


**Figure S4. WGCNA modules identified for aposymbiotic Aiptasia. (a)** Experimental treatments and microbiome traits of aposymbiotic Aiptasia were correlated with module eigengene expression of modules identified through WGCNA. Colours within the heat map correspond to directionality of the Pearson $R^2$ value, in which positive correlations are red and negative correlations are blue. Cells that contain values identify significant relationships between a treatment/trait and a module's eigengene expression. CFU = normalized total Colony Forming Units (**Fig. 1a**); relTotalEndo = relative abundance of the summed counts for Endozoicomonadaceae ASVs (**Fig. 3b**). **(b)** The number of genes belonging to each identified module are identified in a bar graph in which colours correspond to the names of the modules.
